# Muscle spatial lipidomics identifies early ALS signatures in presymptomatic SOD1G93A mice

**DOI:** 10.64898/2026.07.08.737202

**Authors:** Mikel García-Puga, Cristina Huergo, Ainhoa Vidal-Gil, María Rodríguez-Hidalgo, Oihane Pikatza-Menoio, María Levchuk, Amaia Elicegui, Ignacio Azcue, Lucía Romero-Graña, Laura Moreno-Martínez, Rosario Osta, Adolfo López de Munain, José A. Fernández, Sonia Alonso-Martín

## Abstract

Amyotrophic lateral sclerosis (ALS) is a fatal neurodegenerative disease whose diagnosis often remains delayed. Skeletal muscle is increasingly recognized as an early contributor to ALS pathology. Using lipid imaging mass spectrometry (LIMS) in *Tibialis anterior* muscle from *hSOD1^G93A^* mice across disease stages, we identified fiber-type-specific and sex-dependent lipid remodeling. Lipid alterations were detected at the presymptomatic stage, preceding motor neuron loss and clinical symptoms. LIMS distinguished fast-twitch oxidative-glycolytic (type IIA) and glycolytic (type IIB/IIX) fibers and revealed their differential vulnerability to disease. Presymptomatic mutant muscles showed loss of physiological lipid signatures alongside disease-specific lipid changes. Although lipid profiles differed between sexes, ALS-associated alterations enabled accurate discrimination of mutant mice before symptom onset. Importantly, similar disease-related lipid changes were detected in serum, enabling accurate classification of presymptomatic animals. These findings establish lipid remodeling as an early ALS event and highlight novel biomarkers with potential for diagnosis and disease monitoring.

## INTRODUCTION

Amyotrophic lateral sclerosis (ALS) is a progressive and fatal neuromuscular disease characterized by the degeneration of upper and lower motor neurons (MNs), leading to severe skeletal muscle dysfunction. With a global incidence of approximately 1.75 per 100,000 individuals ^1^, most cases are sporadic (sALS), while about 10% are familial (fALS) typically following an autosomal dominant inheritance pattern ^2^. Clinical manifestations include muscle weakness, fasciculation’s, spasticity, impaired fine motor coordination, and occasionally respiratory insufficiency ^3^. The course of the disease is rapid, leading to progressive disability, with a mean survival of 3-5 years from onset, and no effective treatment is currently available. Importantly, ALS is already considered as a non-cell-autonomous disease, meaning that multiple cell types besides MNs, such as glia or skeletal muscle, contribute to the development of the disease ^4–8^.

The first gene linked to ALS was the Cu/Zn superoxide dismutase (SOD1), a cytosolic antioxidant enzyme that converts superoxide radicals into hydrogen peroxide ^9^. Mutations in SOD1 account for approximately 20% of fALS and 2-7% of sALS, with over 230 different mutations identified to date, the majority of which are missense point mutations ^10–13^. The most widely used animal model is the SOD1-G93A mouse, which carries a glycine-to-alanine substitution at position 93 ^14,15^. These mice develop a MN disease characterized by multiple pathological changes reminiscent of human disease, including protein aggregation, oxidative stress, cognitive symptoms, MN loss, motor impairments, and hindlimb paralysis leading to death by 4-months of age ^16–18^. Notably, skeletal muscle alterations occur before MN death and clinical onset, underscoring the peripheral contribution to disease initiation ^19,20^.

The skeletal muscle comprises heterogeneous fiber types with distinct metabolic and contractile properties: slow-twitch oxidative (Type I), fast-twitch oxidative-glycolytic (Type IIA), and fast-twitch glycolytic fibers (Type IIX and IIB) ^21^. In SOD1-G93A mice, fast-twitch fibers are preferentially affected, whereas slow-twitch fibers exhibit greater resistance and become compromised only in later disease stages ^22^. In parallel, compensatory mechanisms such as fiber-type switching occur, leading to a shift toward a more oxidative phenotype ^19,23^.

Parallel to these structural changes in skeletal muscle, metabolic and mitochondrial dysfunction have been studied in ALS. Patients predominantly exhibit hypermetabolism and dyslipidemia, which correlate with symptom severity ^24,25^. In SOD1 transgenic mice, mitochondrial dysfunction includes increased oxidative damage, decreased respiratory activity, swelling, and vacuolization ^26,27^. Importantly, these mitochondrial defects are accompanied by alterations in glycolysis and pyruvate oxidation ^28,29^, as well as disrupted lipid metabolism impairing the utilization of both glucose and fatty acids as energy substrates ^30^. These metabolic deficits are particularly relevant in skeletal muscle fibers, where they may initiate or amplify pathogenic cascades independently of MN loss ^31,32^.

Recent advances in Lipid Imaging Mass Spectrometry (LIMS) offer new opportunities to investigate lipid dysregulation in neurodegenerative diseases (NDD) and other pathologies ^33–36^. Unlike traditional lipidomics, which require tissue homogenization and consequently lose spatial information, LIMS enables direct analysis of lipid species within their native tissue architecture. This technique enables high-resolution mapping of hundreds of individual lipid species across different regions, such as skeletal muscle, and facilitates correlation of lipid signatures with histopathological features in the same tissue section ^33^. Thus, LIMS holds great potential for identifying disease biomarkers for early diagnosis and disease monitoring ^37^.

Biomarkers are essential for early diagnosis, prognosis and treatment in ALS, but their discovery has been challenging partly due to clinical heterogeneity and the lack of validated tissue-specific molecular signatures ^38^. No early and diagnostic biomarkers currently exist for ALS, nor are there validated biomarkers for prognosis or surrogate endpoints to assess treatment efficacy. This diagnostic gap directly delays clinical diagnosis and limits therapeutic options. Increased levels of neurofilament light chain (NfL) are an early indicator of ALS onset and correlate with disease progression ^39,40^. However, NfL lacks disease specificity. Therefore, the discovery and validation of ALS-specific biomarkers are urgently needed to improve diagnosis, facilitate disease monitoring, and enable patient stratification, including the identification of clinical subtypes characterized with distinct disease trajectories and therapeutic responses. Such biomarkers should ideally be measurable in easily accessible biofluids such as serum ^41^. Lipid analysis applied to preclinical models and patient biofluids holds promise for stratifying patients, predicting prognosis, and evaluating response to therapies targeting lipid metabolism ^37,42^.

In this context, we propose using LIMS to study how the lipid profile of skeletal muscle changes in *hSOD1^G93A^* mice during ALS progression at the level of individual fiber types. Our goal is to provide a detailed characterization of lipid remodeling at the cellular level in ALS pathophysiology. Using LIMS, we characterized whole-muscle and fiber-type specific lipid remodeling in skeletal muscle during disease progression. Strikingly, lipid alterations occur already at the presymptomatic stage, prior to MN loss or clinical onset, and differ between fibers. Importantly, spatial lipidomics distinguishes SOD1 mutant from wild-type (WT) animals in both males and females, revealing sex-dependent lipid signatures. We further extended our analysis to serum samples from same animals, identifying circulating lipid species as a new source of biomarkers. Together, our findings demonstrate that fiber-type and sex-dependent lipid alterations emerge early in SOD1 mice, identifying novel candidate tissue and circulating biomarkers with potential utility for early diagnosis, disease monitoring, and patient stratification, an area that remains an unsolved challenge in ALS.

## RESULTS

### Muscle degenerative progression converges with lipid pathway dysregulation in *hSOD1^G93A^* mutant mice

Spinal cord samples from *hSOD1^G93A^* mice and WT littermates were analyzed across presymptomatic, symptomatic, and end-stage phases of ALS progression (**Fig. 1A**). We first characterized spinal cord neurodegeneration in mutant mice using recently developed antibodies against neurofilament light chain (NfL), a sensitive marker of axonal injury and neuronal degeneration ^43^. Following antibody validation (**Fig. S1A**), quantitative analysis revealed an increase in degenerated filaments and neurons (**Fig. S1B**), including MNs, with disease progression. At the presymptomatic stage, the vast majority of filaments and neurons are healthy (**Fig. S1B**), suggesting that at the cellular level, this stage of the disease exhibits low levels of neuronal degeneration. Importantly, a slight increase in degeneration was observed at the symptomatic stage, which became markedly pronounced at the terminal stage (**Fig. S1B**). Calculated of the ratio between degenerated and healthy NfL isoforms demonstrated a notable increase in degeneration at the terminal stage, with no differences at presymptomatic stages (**Fig. 1B**).

**Figure 1.**
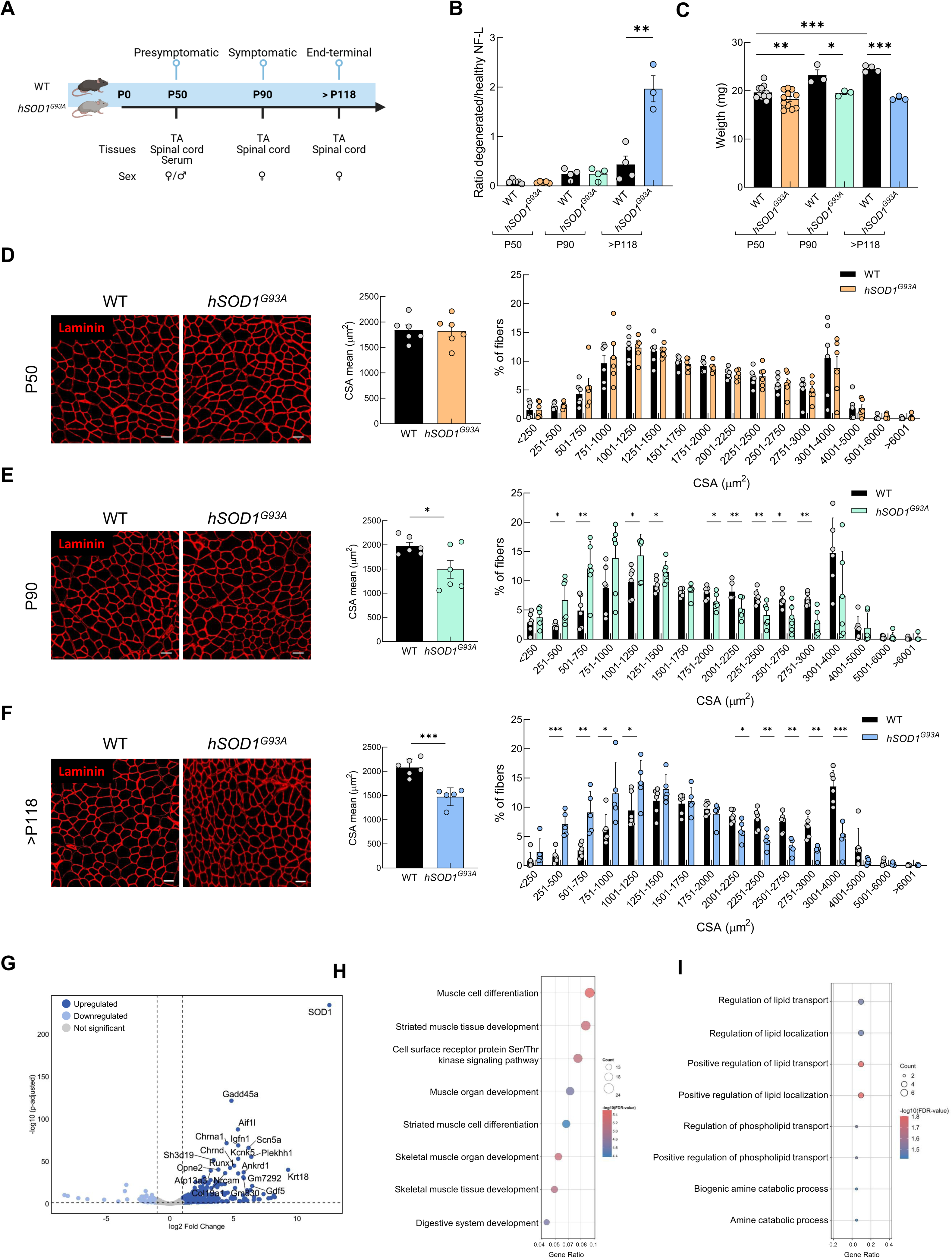
Experimental design and characterization of the skeletal muscle in *hSOD1^G93A^* mice. (**A**) Experimental design and timeline. (**B**) Ratio of fluorescence quantification of CPCA-NfL-Degen (red, marker of the degenerated NfL isoform) to RPCA-NfL-ct (green, marker of healthy NfL isoform) in WT and *hSOD1^G93A^*female mice at different disease stages. (**C**) Body weight of wild-type (WT) and *hSOD1^G93A^* female mice across different disease stages. (**D-F**) Representative images of laminin immunofluorescence, quantification, and frequency distribution of cross-sectional area (CSA) of muscle fibers from WT and *hSOD1^G93A^* female mice at (**D**) presymptomatic P50, (**E**) symptomatic P90, and (**F**) end-stage >P118 disease stages. (**G**) Volcano plot showing statistically significant downregulated (DR, light blue) and upregulated (UR, dark blue) genes in *hSOD1^G93A^*quadriceps from >P118 female mice (fold change ≥ 2 and adjusted *p*-value<0.05). (**H,I**) Dot plots of Gene Ontology (GO) enrichment over-representation analysis for biological processes in (**H**) UR and (**I**) DR genes. Dot size represents the number of genes associated with each GO term, and color indicates the adjusted *p*-value enrichment. P50, P90, and >P118 correspond to postnatal days 50, 90 or >P118, respectively. Bar graphs represent mean ± SEM. N≥5. For normally distributed data with equal variances, one-way ANOVA followed by Tukey’s HSD post hoc test was applied. When variances were unequal, Welch ANOVA followed by Games-Howell post hoc test was used. For non-normally distributed data, the Kruskal-Wallis test followed Wilcoxon rank-sum test was applied. Significant differences are indicated as \**P* < 0.05, ** *P*< 0.01, \*\*\**P* < 0.001.

Then we evaluated changes in body weight over time. WT mice showed a progressive increase in body weight with age (**Fig. 1C**), whereas *hSOD1^G93A^*mice failed to gain weight from the symptomatic stage onward, leading to significantly lower body weight at later stages (**Fig. 1C**). This phenotype is consistent with progressive muscle wasting associated with ALS progression. Histological examination using H&E staining revealed the presence of central-located myonuclei in TA fibers from *hSOD1^G93A^* mutant mice at the terminal stage **(Fig. S2A)**, suggesting tissue impairment, as mature fibers normally exhibit peripheral nuclei. No fat accumulation was detected at any disease stage, with total content remaining below 1% in all conditions **(Fig. S2B)**. However, CSA analysis of muscle fibers across the three different stages of ALS progression indicated muscle atrophy already at the symptomatic stage **(Fig. 1D-F)**. At both symptomatic and terminal stages, mutant mice exhibited a significant reduction of 25% in mean fiber CSA **(Fig. 1E and F**). Moreover, the fiber size distribution in *hSOD1^G93A^* mice was completely altered at both stages of ALS progression **(Fig. 1E and F)**. Mutant mice exhibited a marked shift toward smaller fibers; specifically, the proportion of fibers with a CSA between 250 and 1,000 µm^2^ was approximately 2-fold higher in mutant mice compared to controls. In addition, the population of larger fibers was significantly reduced, with fibers exceeding 2,500-µm^2^ showing an approximate 50% reduction in *hSOD1^G93A^* mutant mice compared to WT mice **(Fig. 1F)**.

Considering that, most pronounced alterations occurred at the terminal stage **(Fig. 1F; Fig. S1A; S2A)**, we therefore performed RNA-seq analysis on quadriceps muscles from WT and *hSOD1^G93A^* mutant mice at this stage to identify metabolic pathways involved in muscle pathology. After confirmation of human *SOD1* overexpression in mutant mice **(Fig. 1G)**, we found 352 genes significantly upregulated (UR) and 92 downregulated (DR) **(Fig. 1G)**. Over-representation analysis (ORA) confirmed that UR genes were enriched in biological processes related to muscle development and muscle homeostasis **(Fig. 1H)**, whereas DR genes were predominantly associated with lipid metabolism-related pathways, such as lipid transport or localization **(Fig. 1I)**. Genes involved in lipid-related pathways included *Map2k6* (also known as *MKK-6)*, a member of the MAP kinase family ^44^, *Eepd1* a nuclease that plays a role in regulation of cholesterol efflux ^45^; *Atp8a1*, a PS flippase in endosomes ^46^; *Abcb4* an ABC transporter that translocate PC across the plasma membrane ^47^; and *Smox* which plays a crucial role in polyamine homeostasis and skeletal muscle differentiation ^48^.

Overall, these results reveal progressive muscle pathology in *hSOD1^G93A^* mice, characterized by dysregulated lipid metabolism and consistent with ongoing muscle degeneration during ALS progression.

### Spatial lipidomic remodeling during postnatal muscle growth

Although ORO staining did not reveal overt changes in total lipid accumulation in TA muscles from *hSOD1^G93A^* mice **(Fig. S2B)**, transcriptomic analyses identified dysregulation of lipid metabolism pathways **(Fig. 1I)**. These findings raised the possibility that ALS progression is accompanied by specific alterations in muscle lipid composition that precede detectable changes in overall lipid storage. To investigate this, we performed untargeted spatial lipidomic profiling of TA muscle across disease stages using LIMS. LIMS is a highly sensitive technique capable of detecting, identifying and relatively quantifying a broad range of lipid species with a spatial resolution of 10 µm/pixel ^36,49^. This approach enables the detection of alterations in membrane-associated and signaling lipids that are not visualized by traditional ORO staining, providing a more comprehensive insight into the lipid remodeling processes underlying ALS pathogenesis.

Applying the workflow described in **Fig. 2A**, we examined TA muscles from WT animals at the three indicated time points. Analysis of the relative abundance of different lipid classes revealed no differences in lipid families from young (P50) to adult stages (P90 or >P118) (**Fig. S3A)**. However, age-dependent alterations in 23 lipid species were detected between young and adult mice **(Fig. 2B-D; Fig. S3)**, and PCA based on these lipids completely separated of P50 control samples from the remaining control groups, indicating a unique lipidomic signature in young animals (**Fig. 2B**). Indeed, P90 and >P118 samples formed overlapping clusters, both clearly distinct from P50 profiles (**Fig. 2B**). Overall, we found that nearly 50% (11 lipids) of the altered lipid species corresponded to PC or PE, while ether- or vinyl ether-linked species derived from PC or PE (PCe/PEe) accounted for 21% (5 lipids) **(Fig. 2C and D; Fig. S3)**. PI represented 13% (3 lipids), whereas PG, PS, SFT, and LSM each contributed 4 % (1 lipid each) **(Fig. 2C and D; Fig. S3)**. The lipid class that experiences the largest remodeling is PC/PE, where a clear reduction in arachidonic acid 20:4 (AA)-containing species (PE 36:4, 37:4, 38:4) occurs, while those containing docosahexaehoic acid 22:6 (DHA) increase, such as PC 36:6/PE 38:6, PC 38:6/PE 40:6 or PC 40:7/PE 42:7 (**Fig. 2C**). Following this trend, an increase in PS 40:6 was observed from P50 to >P118. Likewise, similar changes were observed in PCe/PEe (**Fig. 2D**). Furthermore, image segmentation based on lipidomic profiles, achieved through pixel clustering of similar lipid signatures, uncovered distinct segments corresponding to age-dependent tissue architecture (**Fig. 2E and F)**.

**Figure 2.**
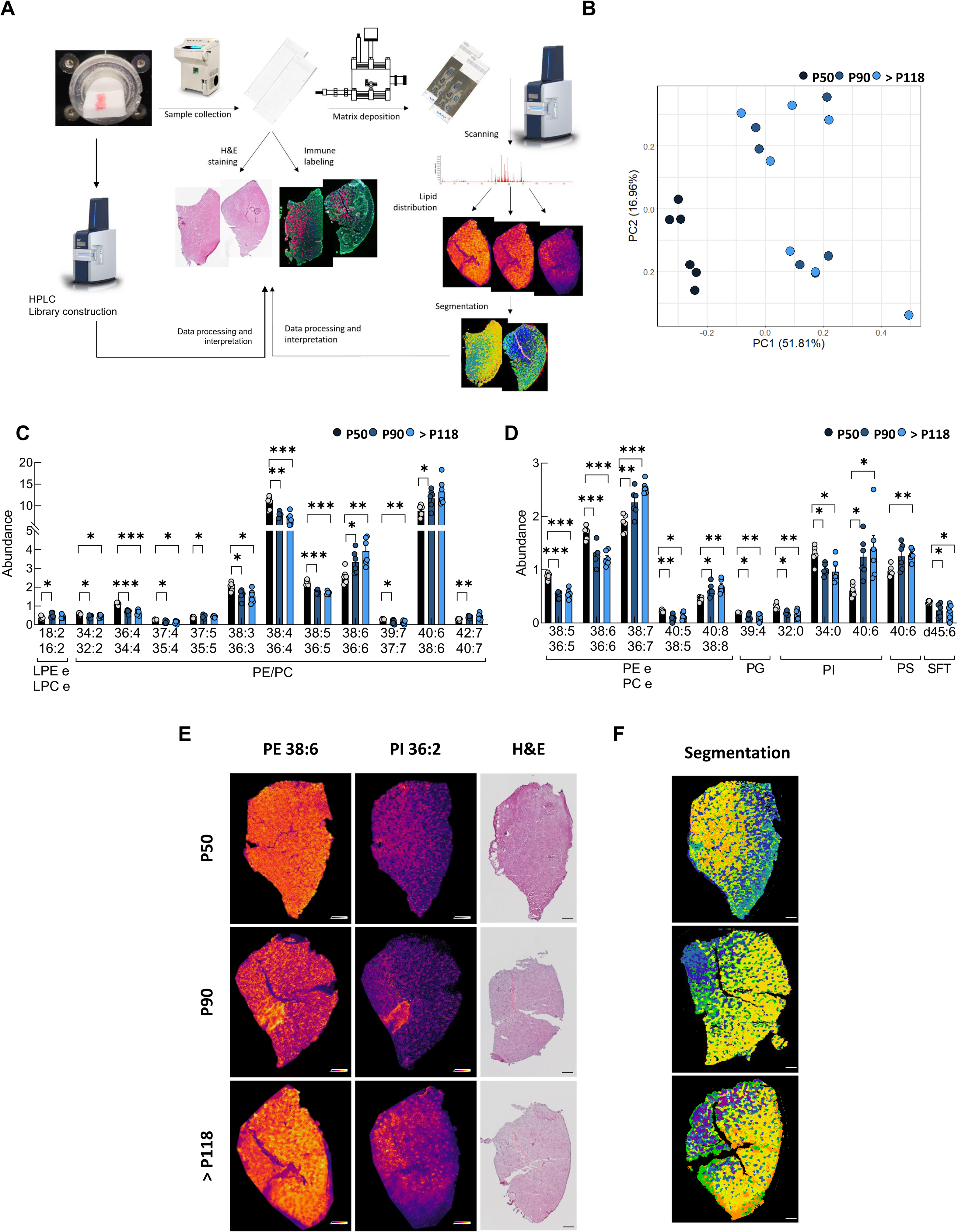
Spatial lipidomic profiling of the skeletal muscle in wild-type control female mice. **(A)** Experimental workflow for untargeted lipid imaging mass spectrometry (LIMS) experiments. **(B)** Principal component analysis (PCA) of significantly altered lipids in the *Tibialis anterior* (TA) muscle of wild-type (WT) female mice at different ages. (**C,D**) Relative abundance of (**C**) LPEe, LPCe, PE, and PC, and (**D**) PEe, PCe, PG, PI, and PS. (**E**) Representative muscle tissue sections showing the spatial distribution of PE 38:6 and PI 36:2 with Hematoxylin and Eosin (H&E) staining. (**F**) Lipid profile-based segmentation. Scale bars: 200 µm. P50, P90 and >P118 correspond to postnatal days 50 (presymptomatic), 90 (symptomatic) or >P118 (end-stage), respectively. Abbreviations: PI, phosphatidylinositol; PE, phosphatidylethanolamine; PC, phosphatidylcholine; PG, phosphatidylglycerol; PS, phosphatidylserine; SFT, sulfatides; LPEe, ether-linked lyso-phosphatidylethanolamine; LPCe, ether-linked lyso-phosphatidylcholine; PEe, ether-linked phosphatidylethanolamine; PCe, ether-linked phosphatidylcholine. Bar graphs represent mean ± SEM. n=6. For normally distributed data with equal variances, one-way ANOVA followed by Tukey’s HSD post hoc test was applied. When variances were unequal, Welch ANOVA followed by Games-Howell post hoc test was used. For non-normally distributed data, the Kruskal-Wallis test followed Wilcoxon rank-sum test was applied. Significant differences are indicated as * *P* < 0.05, ** *P* < 0.01, \*\*\**P* < 0.001.

Together, these findings demonstrate that TA muscles undergo dynamic spatial lipid remodeling throughout postnatal development despite the absence of visible fat accumulation, revealing previously unrecognized age-dependent lipid signatures.

### Disease progression in *hSOD1^G93A^*mutant mice induces spatially resolved lipidomic remodeling

Consistent with control muscle, spatial lipidomic analysis of TA muscles from *hSOD1^G93A^* mice across disease stages revealed the molecular organization of the tissue (**Fig. 3A and B**). Unsupervised lipid-based segmentation identified distinct clusters, which suggests different lipid signatures among muscle fibers, highlighting marked spatial heterogeneity within the muscle. Analysis of the relative abundance of different lipid classes revealed no differences in lipid families in mutant mice from P50 to P90 or >P118 (**Fig. S4A**). However, when we focused the analysis on individual lipid species, we found more pronounced alterations (**Fig. 3C and D; Fig. S4**). PCA of whole-muscle section lipidome in *hSOD1^G93A^* mutant mice showed clustering based not only on age but also on disease progression (**Fig. 3C**). Notably, the *hSOD1^G93A^* lipidome distinguished between the three disease stages; animals at P50, P90, and >P118 clustered separately from each other (**Fig. 3C**), with 24 lipid species altered mice throughout disease progression (**Fig. 3D; Fig. S4**). Among these, 45% (11 lipids) corresponded to PC or PE, followed by PCe/PEe (12%, 3 lipids), PS, PI and SM (8%, 2 lipids each), among other low-abundance lipid species (**Fig. 3D; Fig. S3**).

**Figure 3.**
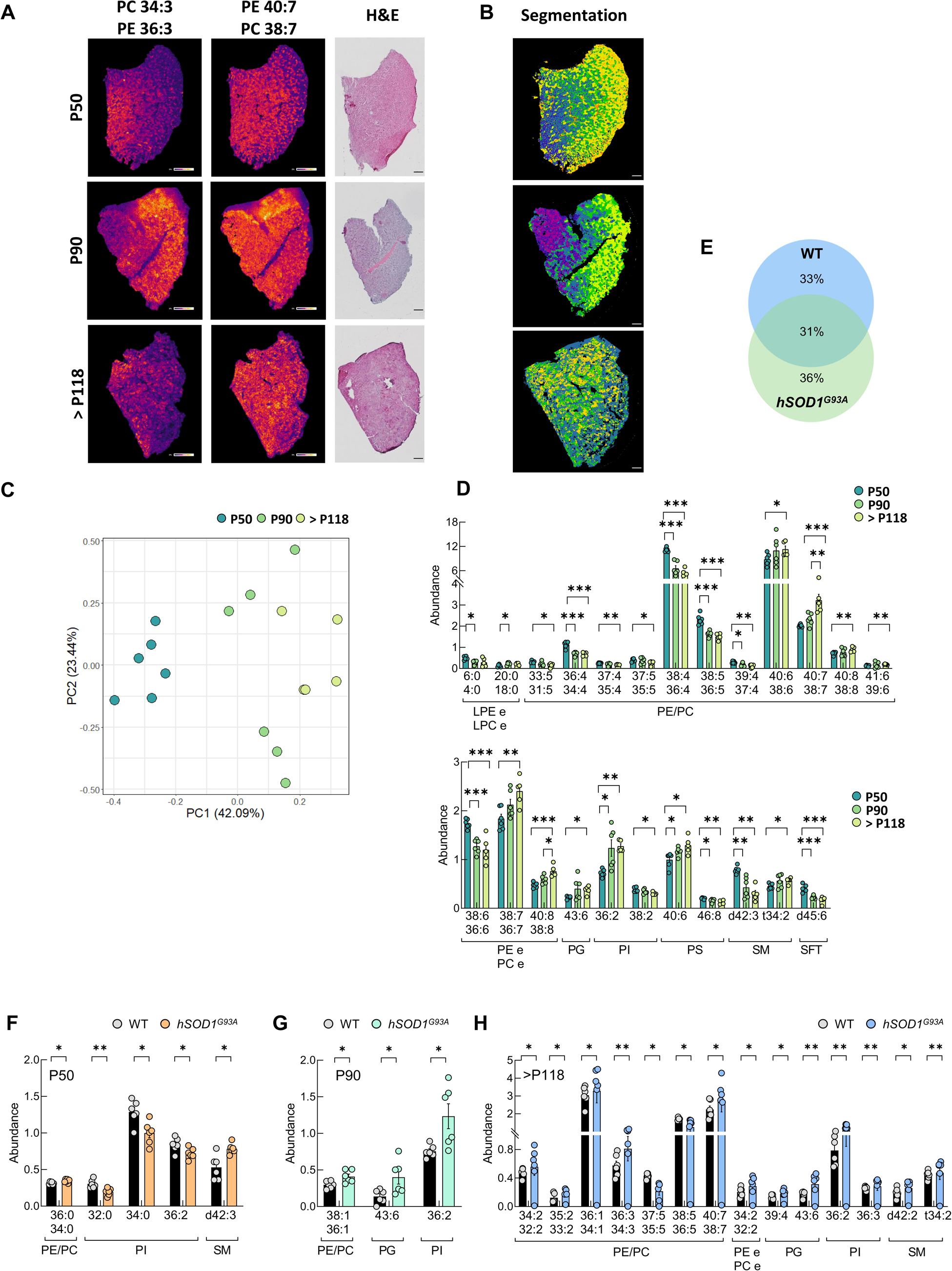
Altered spatial lipidome in the skeletal muscle of *hSOD1^G93A^* mice. (**A**) Representative *Tibialis anterior* (TA) muscle sections showing distribution of PC 34:3 and PE 40:7 with Hematoxylin and Eosin (H&E) staining. (**B**) Lipid profile-based segmentation of TA muscle. Scale bars: 200 µm. (**C**) Principal component analysis (PCA) of significantly altered lipids in *hSOD1^G93A^* female mice at different ages. (**D**) Relative abundance of LPEe, LPCe, PE, PC, PEe, PCe, PG, PI, PS, SM, and SFT. (**E**) Venn diagram identifying the number of differentially altered lipid species unique to and shared between wild-type (WT) control and *hSOD1^G93A^*mice. (**F-H**) Relative abundance of different lipid species at (**F**) P50, (**G**) P90, and (**H**) >P118. P50, P90 and >P118, correspond to postnatal days 50 (presymptomatic), 90 (symptomatic) or >P118 (end-stage), respectively. Abbreviations: PI, phosphatidylinositol; PE, phosphatidylethanolamine; PC, phosphatidylcholine; PG, phosphatidylglycerol; PS, phosphatidylserine; SM, sphingomyelin; SFT, sulfatides; LPEe, ether-linked lyso-phosphatidylethanolamine; LPCe, ether-linked lyso-phosphatidylcholine; PEe, ether-linked phosphatidylethanolamine; PCe, ether-linked phosphatidylcholine. Bar graphs represent mean ± SEM. n≥5. For normally distributed data with equal variances, one-way ANOVA followed by Tukey’s HSD post hoc test was applied. When variances were unequal, Welch ANOVA followed by Games-Howell post hoc test was used. For non-normally distributed data, the Kruskal-Wallis test followed Wilcoxon rank-sum test was applied. Significant differences are indicated as \**P* < 0.05, \*\**P* < 0.01, \*\*\**P* < 0.001.

Furthermore, comparison of altered lipid species between control and mutant mice revealed that 36% were shared between genotypes, likely reflecting age-related lipid remodeling independent of disease. In contrast, 33% were uniquely altered in WT mice, suggesting adaptive lipid changes that fail to occur in mutant mice, whereas 36% were exclusive to *hSOD1^G93A^*mice, indicating distinct lipid remodeling trajectories during ALS progression **(Fig. 3E)**. In summary, despite unchanged global lipid class abundance, the lipidome of *hSOD1^G93A^* mice undergoes progressive remodeling at the species level throughout ALS progression. The clear segregation of presymptomatic from symptomatic and terminal stages, together with the identified 24 altered lipid species, reveals a dynamic lipid fingerprint that correlates with disease progression in mutant muscles.

Finally, examination of the differences between mutant and control mice at three disease stages showed significant changes in 5 lipid species at P50, 3 species at P90, and a marked increase in lipid changes at the terminal stage, with changes in 14 lipid species (**Fig. 3F-H; Fig. S5**).

Collectively, these results demonstrate that a dynamic lipid fingerprint in skeletal muscle characterizes ALS progression in the SOD1 mouse model, with the most extensive alterations occurring at the terminal stage. Thus, the ability of the lipidome to distinguish between presymptomatic, symptomatic and terminal stages support the potential use of specific lipid species as disease progression biomarkers.

### Fiber-type-specific lipid remodeling accompanies disease progression in SOD1 mutant mice

In rodents, TA muscles are predominantly composed of type IIA (fast-twitch oxidative-glycolytic) and type IIB fibers, along with type IIX fibers (both fast-twitch glycolytic) ^21^. Initial analysis revealed that TA is composed exclusively of type II fast fibers, with no significant changes in their relative proportion across disease stages **(Fig. S6A)**. However, detailed analysis demonstrated that type IIB fibers exhibited a 30% and 40% reduction in CSA at symptomatic and terminal stages, respectively, with no changes observed in type IIA or IIX fibers **(Fig. S6B)**. Distribution analysis of type IIA and IIX fibers exhibited fewer morphological changes than type IIB fibers at both symptomatic and terminal stages **(Fig. S6C-E)**. Type IIB fibers in *hSOD1^G93A^* mice further revealed a shift, showing 8-fold and 4-fold increase in smaller fibers (251-1,500 µm^2^ CSA) at symptomatic and terminal stages, respectively **(Fig. S6C-E)**. In addition, larger fibers with a CSA of 2,501-3,000 µm^2^ showed a 60% and 70% reduction at symptomatic and terminal stages, respectively **(Fig. S6D)**.

To verify whether muscle fiber identity is reflected in the spatial lipidome, we performed fiber-resolved lipidomic analysis in WT mice. Remarkably, distinct lipid signatures accurately discriminated the different fiber types and closely matched the classification obtained by immunofluorescence, with a minimal set of only three lipids (PE 38:4/PC 36:4; PE 40:6/PC 38:6 and PI 34:0) being sufficient to achieve accurate fiber-type identification (**Fig. 4A and B**). Each lipidome-based cluster could be linked to a specific fiber type, demonstrating specificity of the lipid fingerprint (**Fig. 4A and B**). Thus, we investigated the specific differences in lipid composition among different fiber types. At P50, IIA, IIB, and IIX fibers exhibited distinct lipid profiles in WT mice (**Fig. 4C**), with PG being the lipid class that experiences the largest change between IIA and IIB fibers (**Fig. S7A**). At the individual lipid species level, we identified changes in 32 lipid species, most of them belonging to PC/PE (53%), followed by PI (22%) and SM (12.5%), along with PS and SFT (6% each) (**Fig. S7**). We then assessed lipidomic changes across the three fiber types during postnatal development in WT mice. As before, P50 mice were clearly distinct from the other age groups across all three-fiber types analyzed. In contrast, the lipidomic profiles did not discriminate between P90 and >P118 mice in any fiber type (**Fig. 4D; S8-S10**), suggesting that lipid remodeling largely occurs during early postnatal development and reaches a stable state in adulthood.

**Figure 4.**
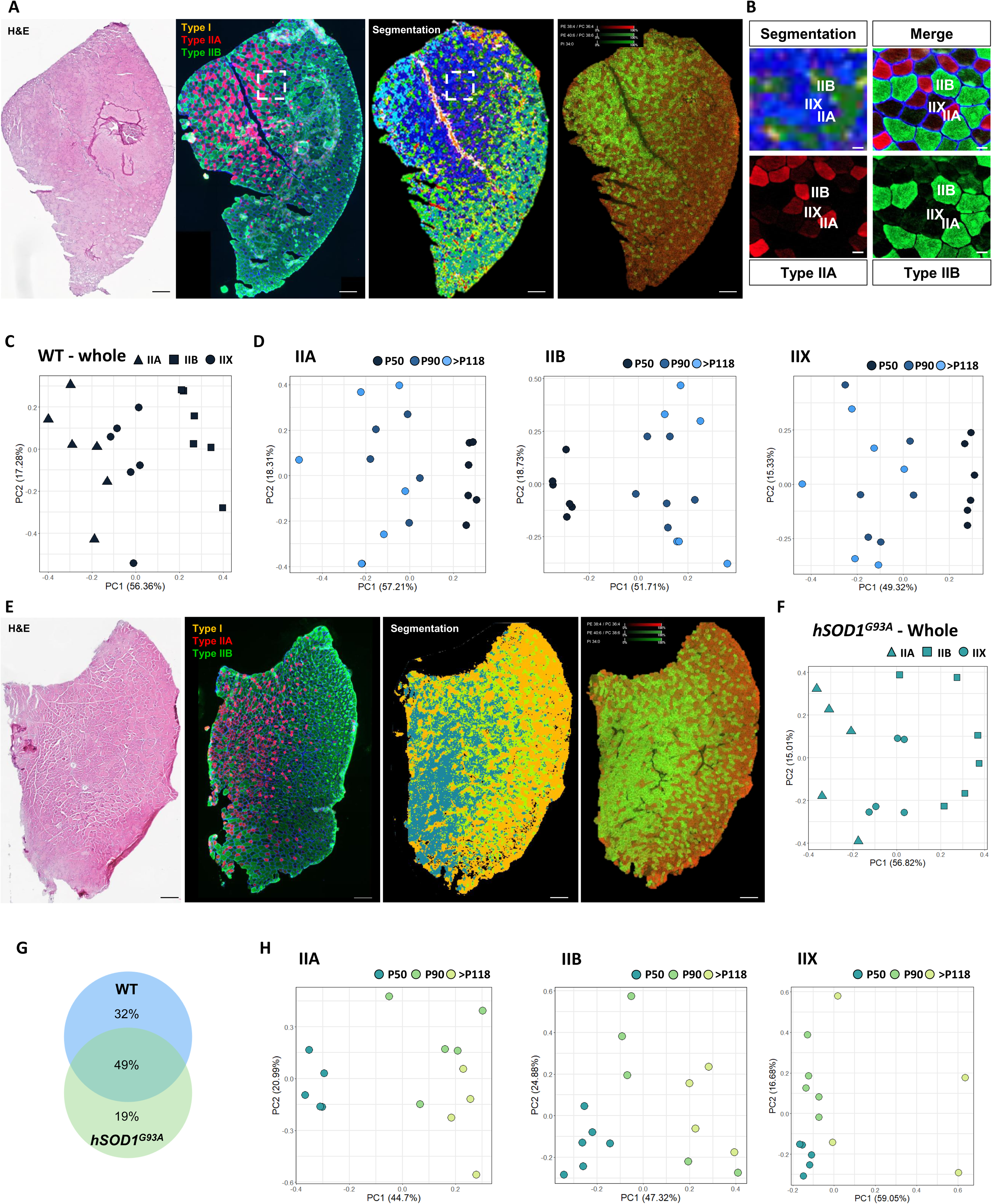
Fiber type-specific lipidomics in the skeletal muscle in *hSOD1^G93A^* mice. (**A**) Representative tissue sections of Hematoxylin and Eosin (H&E) staining, fiber type (I, IIA, IIB, IIX; with IIX identified as the unstained) immunofluorescence, lipid profile-based segmentation and lipid distribution of PE 38:4/PC 36:4; PE 40:6/PC 38:6 and PI 34:0 in wild-type (WT) female control mice. Red squared is enlarged in B). Scale bars: 200 µm. (**B**) Lipid profile-based segmentation and fiber type immunostaining from (**A**). Scale bar: 20 µm. (**C**) Principal component analysis (PCA) of significantly altered lipid species from type IIA, IIB and IIX fibers in WT at P50. (**D**) PCA of significantly altered lipid species from specific type IIA, IIB, or IIX fibers in WT mice at P50, P90, and >P118. (**E**) Representative tissue sections of H&E staining, fiber type (I, IIA, IIB, IIX; with IIX identified as the unstained) immunofluorescence, lipid profile-based segmentation and lipid distribution of PE 38:4/PC 36:4; PE 40:6/PC 38:6 and PI 34:0 in *hSOD1^G93A^* (SOD1) female mice. Scale bars: 200 µm. (**F**) PCA of dysregulated lipid species from IIA, IIB and IIX fibers in SOD1 at P50. (**G**) Venn diagram showing the number of differentially altered lipid species from panels (**C**) and (**F**). The diagram illustrates the overlap between WT and SOD1 female mice. (**H**) PCA plots of dysregulated lipid species from specific IIA, IIB, or IIX fibers in SOD1 mice during disease progression. N≥4. P50, P90 and >P118 correspond to postnatal days 50 (presymptomatic for SOD1), 90 (symptomatic for SOD1) or >P118 (end-stage for SOD1), respectively. For normally distributed data with equal variances, one-way ANOVA followed by Tukey’s HSD post hoc test was applied. When variances were unequal, Welch ANOVA followed by Games-Howell post hoc test was used. For non-normally distributed data, the Kruskal-Wallis test followed Wilcoxon rank-sum test was applied. Significant differences are indicated as \**P* < 0.05, \*\**P* < 0.01, \*\*\**P* < 0.001.

To assess disease-related alterations, we examined fiber-type-specific lipidomic changes during ALS progression. As in WT, lipidome-based segmentation accurately recapitulated the distinct fiber types (IIA, IIB, and IIX) of presymptomatic *hSOD1^G93A^* mice (**Fig. 4E**). At this stage, the three fiber types exhibited clearly distinct lipid profiles, with a total of 33 significantly altered lipid species (**Fig. 4F; Fig. S11**). To determine whether these lipid signatures were specific to the mutant genotype or preserved under physiological conditions, we compared the lipid species that differed among IIA, IIB, and IIX between WT and mutant mice at the presymptomatic stage. Strikingly, 49% of the differentially altered lipid species were shared between genotypes, suggesting a common lipid-based signature of fiber-type specification. Furthermore, 32% were unique to WT mice, suggesting that mutants lose a substantial portion of the physiological lipid program, and 19% were exclusive to *hSOD1^G93A^* mice, revealing early pathological lipid remodeling (**Fig. 4G**). We finally analyzed lipid remodeling across disease progression in type IIA, IIB, and IIX fibers from *hSOD1^G93A^*mice, identifying disease-dependent fiber type-specific lipid remodeling trajectories in each fiber type. Notably, lipid profiles clearly discriminated presymptomatic, symptomatic, and end-stage animals, revealing progressive remodeling from disease onset to terminal stages (**Fig. 4H; Fig. S12-S14**).

In summary, these results reveal a previously unrecognized fiber-specific lipid signature in TA muscle that distinguishes fast-twitch oxidative-glycolytic (IIA) from fast-twitch glycolytic (IIB/IIX) fibers. Mutant-specific alterations reveal early pathological lipid remodeling that progresses across ALS stages.

### Presymptomatic lipidomics identifies sex-specific signatures with diagnostic and prognostic potential

Given the well-established sex differences in ALS, with males exhibiting higher disease incidence and faster disease progression than females ^50^; we extended our spatial lipidomic analysis to presymptomatic male *hSOD1^G93A^* mice to complement our findings in females. We first measured body weight and quantified healthy and degenerated filaments, neurons, and MNs (**Fig. 5A and B**), detecting no differences between genotypes. Consistent with this, TA muscle histology showed normal morphology with no central nucleated fibers and residual neutral lipid accumulation (<1%; **Fig. 5C**). Moreover, mean fiber size, fiber size distribution, and fiber type composition (IIA, IIB and IIX fibers) were comparable between groups (**Fig. 5C and D**). Together these findings indicate that male *hSOD1^G93A^* mice at P50 remain presymptomatic, displaying no detectable neuromuscular pathology and MN degeneration, consistent with our observations in females. Analysis of lipid class relative abundance revealed significant alterations in several lipid families, including SFT, PI, and lyso- phospholipids derived from PC, PE, and SFT (**Fig. S15A)**. At the individual species level, 25 lipids were differentially affected in *hSOD1^G93A^* mice, with PE, PC, SM, and lyso- phospholipids showing the most lipid prominent changes (**Fig. 5E and F; Fig. S15**).

**Figure 5.**
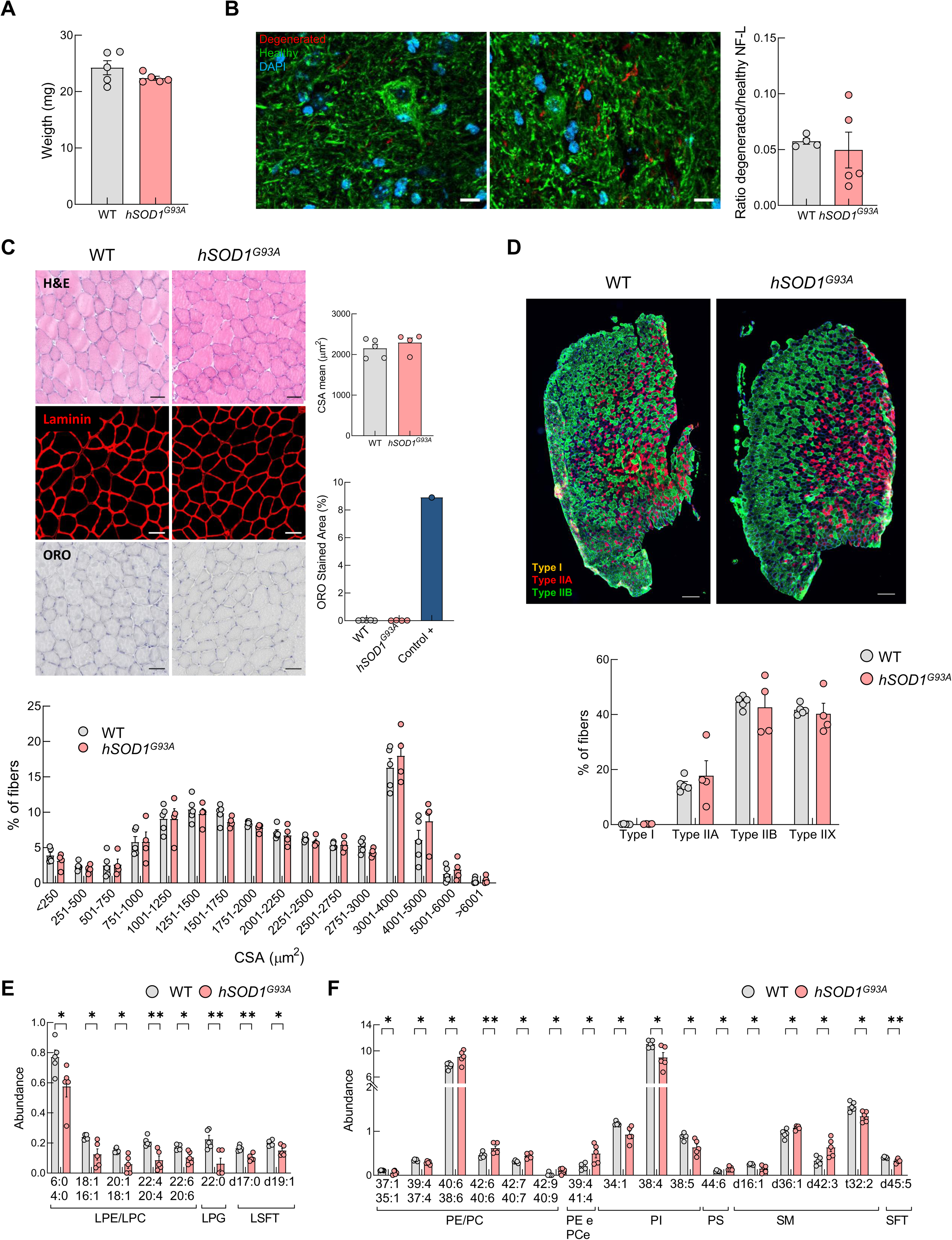
Skeletal muscle characterization and lipidomics in presymptomatic *hSOD1^G93A^* male mice. (**A**) Body weight of wild-type (WT) and *hSOD1^G93A^* male mice at P50. (**B**) Representative images and ratio of fluorescence quantification of CPCA-NfL-Degen (red, marker of the degenerated NfL isoform) to RPCA-NfL-ct (green, marker of healthy NfL isoform) in WT and *hSOD1^G93A^*male mice at P50. Scale bar = 10 µm. (**C**) Representative images of the *Tibialis anterior* (TA) muscle from WT and *hSOD1^G93A^* male mice at P50 stained with Hematoxylin and Eosin (H&E), laminin immunofluorescence, quantification, and frequency distribution of cross-sectional area (CSA), and Oil Red O (ORO) with quantification. Mouse skin tissue was used as a positive control for ORO staining. Scale bars = 50 µm. (**D**) Representative tissue section of fiber type immunofluorescence and quantification of type I, IIA, IIB, and IIX fibers; with IIX identified as the unstained. Scale bars = 200 µm. (**E,F**) Relative abundance of (**E**) LPE, LPC, LPG, and LSM, and (**F**) PE, PC, PEe, PCe, PI, PS, SM, and SFT in WT and *hSOD1^G93A^* male mice at P50. P50 corresponds to postnatal day 50 (presymptomatic for *hSOD1^G93A^*). Abbreviations: PI, phosphatidylinositol; PE, phosphatidylethanolamine; PC, phosphatidylcholine; PS, phosphatidylserine; SM, sphingomyelin; SFT, sulfatides; LPE, lyso-phosphatidylethanolamine; LPC, lyso-phosphatidylcholine; LPG, lyso-phosphatidylglycerol; LST, lyso sulfatides; PEe, ether-linked phosphatidylethanolamine; PCe, ether-linked phosphatidylcholine;. Bar graphs represent mean ± SEM. N=5. For normally distributed data with equal variances, one-way ANOVA followed by Tukey’s HSD post hoc test was applied. When variances were unequal, Welch ANOVA followed by Games-Howell post hoc test was used. For non-normally distributed data, the Kruskal-Wallis test followed Wilcoxon rank-sum test was applied. Significant differences are indicated as \**P* < 0.05, \*\**P* < 0.01, \*\*\**P* < 0.001.

In summary, male *hSOD1^G93A^* mice at P50 remain presymptomatic, mirroring the status observed in females. Nevertheless, whole-muscle lipidomics revealed significant genotype-dependent alterations, indicating that lipid remodeling precedes overt neuromuscular pathology in ALS.

### Presymptomatic lipidomics signatures in muscle and serum support ALS diagnosis and prognosis

Having validated spatial lipidomics in ALS muscle and confirmed the presymptomatic status of both male and female *hSOD1^G93A^* mice at P50, we next sought to identify early lipidomic signatures associated with disease. Such presymptomatic alterations may provide a basis for biomarker development and facilitate prediction of potential disease progression. We first analyzed whole-muscle and fiber-specific lipidomes in presymptomatic female mice, revealing distinct lipid signatures that clearly distinguish *hSOD1^G93A^*mice from WT controls (**Fig. 6A-D**). Despite the relatively limited number of altered lipid species compared with later disease stages, these differences robustly segregated mutant and control animals (**Fig. S16-S18**). We then developed classification models to assess the biomarker potential of candidate lipid species. Across both whole-muscle and fiber-specific panels, all models achieved high AUC, sensitivity, and specificity values, supporting the utility of presymptomatic lipid signatures for ALS diagnosis (**Fig. 6A-D**). We next extended the analysis to male mice. Spatial lipidomics robustly separated between *hSOD1^G93A^*and control animals based on both whole-muscle and fiber-type-specific lipidomes, and classification models achieved again high predictive performance across all lipid panels (**Fig. 6E-H; Fig. S19-S21**). Strikingly, the altered lipid signatures showed minimal overlap between sexes. Indeed, only a single lipid specie was shared in the whole lipidome analysis, whereas all other comparisons identified distinct male- and female-specific lipid panels (**Fig. 6I**).

**Figure 6.**
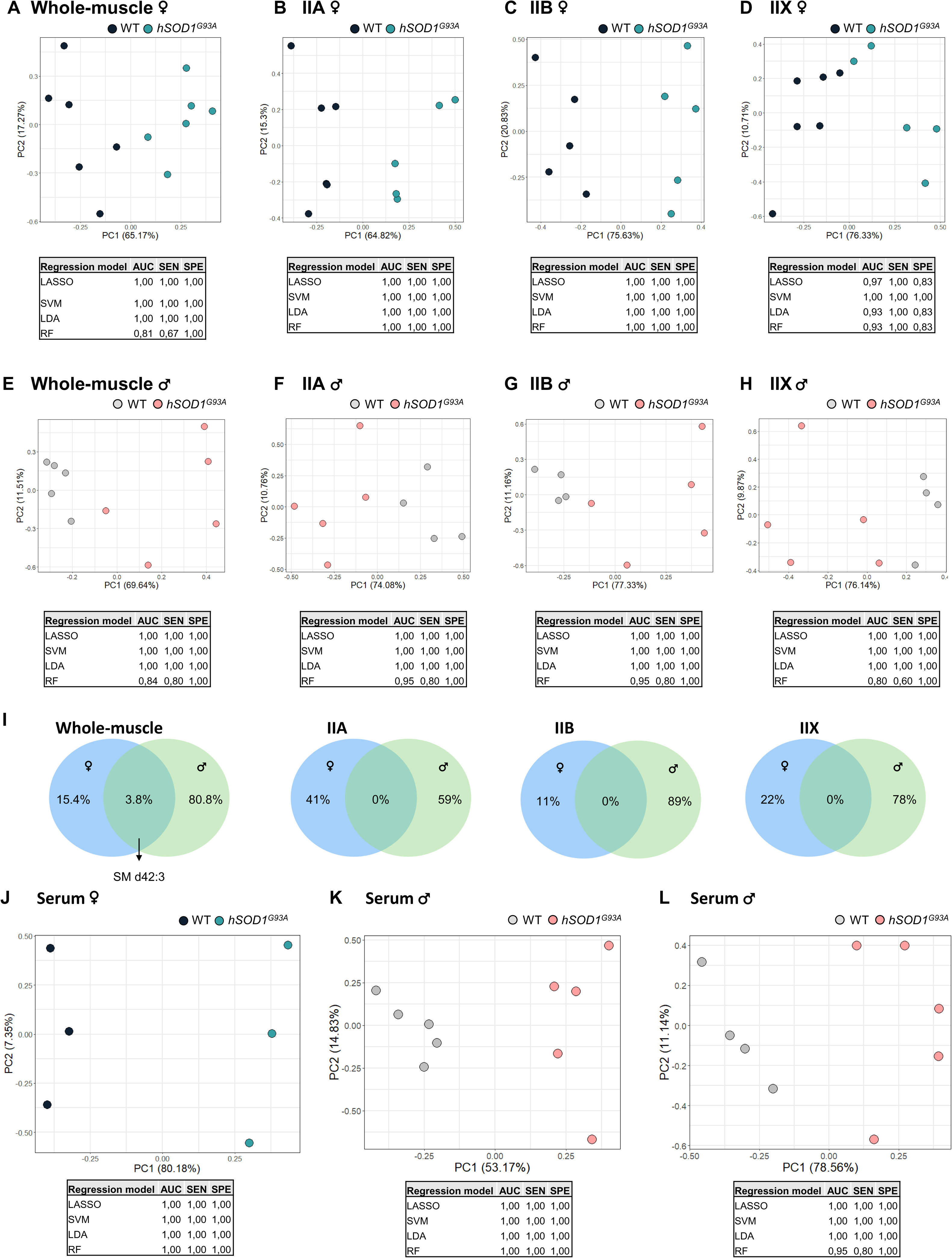
Whole-section, fiber type-specific and serum lipidomic signatures in *hSOD1^G93A^* mice at presymptomatic stage. (**A-H**) Principal component analysis (PCA) plots and regression models metrics of significantly altered lipid species from whole muscle sections and individual fiber type-specific profiles (IIA, IIB, and IIX) at the P50 stage in (**A-D**) female or (**E-H**) male *hSOD1^G93A^*(SOD1) mice compared with wild-type (WT) control mice. (**I)** Venn diagram showing the number of differentially altered lipid species in whole muscle sections and individual fiber types between male and female SOD1 mice compared with WT controls. (**J,K**) PCA plots and regression models metrics of significantly altered lipid species from serum of (**J**) female (n=3) and (**K**) male (n=5) SOD1 mice compared with WT controls. (**L**) PCA plot of a reduced 4-lipid panel shared between muscle and serum, shown in male serum. P50 corresponds to postnatal day 50 (presymptomatic for SOD1). N=5.

Given the strong discriminative power of presymptomatic muscle lipid signatures, we next asked whether similar lipid alterations could be detected in serum. Given that serum can be obtained through minimally invasive procedures, it represents an attractive biofluid for biomarker discovery and longitudinal disease monitoring. Analysis of presymptomatic female serum identified 26 altered lipids that clearly distinguished *hSOD1^G93A^*mice from WT controls (**Fig. 6J**). Applying the same approach to males revealed 41 differentially altered lipids, which likewise enabled complete separation of mutant and WT mice (**Fig. 6K**). Comparison of the male serum and the whole-muscle lipid panels identified 4 lipids that were significantly altered in both tissues (**Fig. 6A; Fig. S16**). Remarkably, this minimal 4-lipid signature was sufficient to accurately classify presymptomatic mice (**Fig. 6L**).

Together, these findings reveal that sex-dependent lipidomic changes arise before overt disease manifestation, establishing lipid remodeling as an early event in ALS pathogenesis. The robust classification performance observed in both muscle and serum supports the potential of these signatures for early diagnosis, patient stratification, and disease monitoring.

## DISCUSSION

ALS remains a rapidly progressive NDD for which reliable biomarkers and effective therapies are still lacking. Although traditionally viewed as a MN disorder, increasing evidence supports an active contribution of peripheral tissues, including skeletal muscle, to disease onset and progression ^8,51^. Using high-resolution spatial lipidomics in *hSOD1^G93A^* mice, we identify lipid remodeling as an early feature of ALS pathogenesis that emerges before overt neuromuscular pathology. Our results reveal extensive fiber type-specific and sex-dependent lipid alterations and identify circulating lipid signatures capable of discriminating presymptomatic animals. Together, these findings establish lipid metabolism as an early and previously underappreciated component of ALS pathology.

A major finding of this study is that lipid remodeling is already evident at the presymptomatic stage, preceding detectable muscle atrophy and neurodegeneration. These observations further support the concept that skeletal muscle is not merely a passive target of MN dysfunction but an active participant in disease progression. Previous studies have demonstrated that muscle-specific expression of mutant SOD1 is sufficient to induce pathological alterations ^52,53^, and that neuromuscular junction dysfunction and muscle atrophy occur before MN loss in *hSOD1^G93A^*mice ^54,55^. Our results extend these observations by identifying metabolic remodeling of muscle lipids as an additional early event associated with disease development, suggesting that disruption of lipid homeostasis may contribute to the earliest stages of ALS pathology.

The preferential vulnerability of type IIB fast-twitch glycolytic fibers observed in our study is consistent with previous reports in ALS patients and experimental models ^56–58^. While the mechanisms underlying this selective sensitivity remain unclear, the greater metabolic demand and lower oxidative capacity of glycolytic fibers may contribute to their increased susceptibility ^21,59^. Importantly, spatial lipidomics revealed that each muscle fiber type possesses a unique lipid signature and that these physiological signatures are profoundly altered in presymptomatic *hSOD1^G93A^* mice. The emergence of mutation-specific lipid profiles before overt pathology suggests that fiber-type-specific lipid composition may influence disease susceptibility and provides a potential metabolic basis for selective fiber degeneration in ALS.

Although lipid dysregulation is increasingly recognized as a hallmark of neurodegeneration ^60,61^, most studies have focused on the central nervous system (CNS). In skeletal muscle, previous work has described an early shift towards lipid utilization together with increased lipolysis during disease progression ^62–64^. Our findings extend these observations by providing a spatially resolved characterization of muscle lipid remodeling. In particular, glycerophospholipids, especially PC and PE species, emerged as major targets of early dysregulation. Given their central role in membrane organization, signaling, and autophagy, alterations in these lipid classes may have broad consequences for muscle homeostasis. Consistent transcriptomic changes in genes involved in phospholipid transport, membrane remodeling, and muscle maintenance further support disruption of membrane lipid homeostasis. Moreover, alterations in ether-linked phospholipids suggest impaired plasmalogen metabolism, potentially increasing vulnerability to oxidative stress, a well-established feature of ALS muscle pathology.

Importantly, our observations are not restricted to the SOD1 model. Lipid dyshomeostasis has also been described in *TDP-43^Q^*^331^*^K^* mice, in which skeletal muscle exhibits substantial metabolic alterations, in some cases exceeding those observed in the spinal cord ^65^. Together with previous reports in ALS patients, these findings suggest that lipid remodeling may represent a common pathological feature across genetically distinct forms of ALS and further support skeletal muscle as an important site of metabolic dysfunction.

Another striking observation was the marked sexual dimorphism detected at the presymptomatic stage. Male and female *hSOD1^G93A^* mice displayed largely distinct lipid signatures, sharing only one altered lipid species in whole-muscle analyses. These findings are consistent with the known sex differences in ALS incidence, progression, and metabolism ^50,66^. Our results extend these observations by demonstrating that sex-dependent metabolic remodeling is already evident before symptom onset, suggesting that intrinsic metabolic differences may contribute to disease susceptibility and progression. These findings highlight the importance of incorporating sex as a biological variable in both biomarker development and therapeutic studies.

Importantly, lipid alterations were not restricted to skeletal muscle. Analysis of presymptomatic serum identified lipid signatures that robustly discriminated mutant from control animals. Remarkably, a minimal four-lipid panel shared between muscle and serum accurately classified presymptomatic mice. These observations are consistent with emerging evidence of altered lipid metabolism in ALS patient plasma ^67,68^ and suggest that circulating lipids may reflect ongoing pathological processes occurring within skeletal muscle. As such, lipidomic biomarkers may provide a minimally invasive approach for early diagnosis, patient stratification, and disease monitoring.

A major strength of this work is the application of high-resolution spatial lipidomics, which preserves tissue architecture while enabling lipid profiling at the level of individual muscle fibers. By integrating disease-stage, fiber-type, and sex-specific analyses, this approach reveals biological information that would be inaccessible using conventional bulk lipidomic methods. Future studies will be required to determine the extent to which these signatures are conserved across human ALS subtypes and whether they can be leveraged for clinical biomarker development.

Finally, our findings place lipid metabolism among the earliest alterations associated with ALS pathogenesis and further support the growing interest in metabolic interventions as therapeutic strategies. Both preclinical and clinical studies indicate that manipulation of lipid metabolism can influence disease progression, supporting the concept that lipid pathways may represent tractable therapeutic targets. In summary, our study identifies extensive lipid remodeling as an early and previously underappreciated feature of ALS pathogenesis. Using spatial lipidomics, we uncover fiber type-specific and sex-dependent lipid signatures that emerge before overt neuromuscular pathology and extend into the circulation. These findings establish skeletal muscle as an important site of early metabolic dysfunction and highlight lipidomic signatures as promising candidates for diagnosis, stratification, and disease monitoring in ALS.

## METHODS

### Animals and sample collection

The *hSOD1^G93A^* transgenic mouse strain (*B6SJL-Tg[SOD1-G93A]1Gur*; Strain #002726, RRID:IMSR_JAX:002726), which carries a high copy number of the mutant human SOD1 transgene, was used in this study. Non-transgenic B6SJL wild-type (WT) littermates were used as controls. *hSOD1^G93A^* mice were obtained from The Jackson Laboratories (Bar Harbor, ME, USA). Genotyping of hemizygous transgenic and WT offspring was performed by polymerase chain reaction amplification of genomic DNA isolated from tail tissue. Mice were housed in ventilated cages under a 12-hour (h) light-dark cycle with *ad libitum* access to food and water. Based on disease progression, the presymptomatic stage was defined as postnatal day 50 (P50), the symptomatic stage as P90, and the terminal stage as the point when mice were unable to right themselves within 30 seconds when placed supine (>P118). Mice were euthanized at the indicated stages. We used female *hSOD1^G93A^* mice and their WT littermates at P50 (n=6 per group), P90 (n=6 per group), and P118 (WT, n=6; *hSOD1^G93A^* n=5), as well as male mice at P50 (n=5 per group). *Tibialis anterior* (TA) muscles were dissected, snap-frozen in isopentane pre-chilled with liquid nitrogen and stored at -80 °C, as previously described ^69,70^. Spinal cords were dissected and frozen following the same procedure as the TA, but cryopreserved in Tissue-Tek® O.C.T. Compound (4583, Sakura, Torrance, CA, USA). All animal handling and protocols were approved by the Animal Care Ethics Committee of Biogipuzkoa Institute and the Provincial Council of Gipuzkoa (PRO-AE-SS-193 [OH-20-37] and PRO-AE-SS-235 [OH21-42]), the Ethics Committee for Animal Experiments of the Universidad de Zaragoza (PI52/15). All procedures were conducted in conformity with EU guidelines and regulations for animal welfare (3R principles).

### Histological staining

14-µm thick fresh-frozen cryosections were obtained and fixed in 4% paraformaldehyde (PFA, 15713, Electron Microscopy Sciences, Morgantown, PA, USA) for 10 minutes (min). Following fixation, sections were processed for Hematoxylin & Eosin (H&E) staining. Briefly, sections were washed twice in phosphate-buffered saline (PBS, 10 min each), stained with Harris hematoxylin (253949, Panreac AppliChem, Darmstadt, Germany) for 1 min, rinsed in water, counterstained with eosin (251301, Panreac AppliChem) for 30 seconds (s), and rinsed again in water. Subsequently, sections were dehydrated through a series of ethanol washes: 10 times in increasing ethanol concentrations (30%, 50%, 70%, 85%, and 95%), followed by two 20-min incubations in 100% ethanol. Finally, sections were cleared by incubating twice in xylene for 30 min each, and mounted with DPX mounting medium (06522, Sigma-Aldrich, St. Louis, MO, USA). For Oil Red O (ORO) staining, tissue sections were washed twice in distilled water (dH_2_O) for 10 min each, then dehydrated with 100% 1,2-propanediol (98039, Sigma-Aldrich, St. Louis, MO, USA) for 5 min prior to incubation with 0.5% ORO solution (O0625, Sigma Aldrich, St. Louis, MO, USA) for 10 min. Sections were then washed twice in 85% 1,2-propanediol (398039, Sigma-Aldrich, St. Louis, MO, USA) and twice in dH_2_O for 10 min each. Nuclei were counterstained with Harris hematoxylin (253949, Panreac AppliChem, Darmstadt, Germany) for 1 min, washed twice in dH_2_O and mounted with Fluoromount-G (0100-01, Southern Biotech, Birmingham, AL, USA).

### Immunofluorescence

Cryosections of fresh-frozen tissue (14 µm-thick TA muscles and 20 µm-thick spinal cords) were used. While muscles sections were fixed and washed as for histological staining, spinal cord sections were fixed in cold 4% PFA for 15 min, and then washed three times in PBS for 10 min each. Blocking and permeabilization were performed using a solution containing 4% bovine serum albumin (BSA, 001-000-162, Jackson ImmunoResearch Inc., West Grove, PA, USA), 1% goat serum (GS, 005-000-121, Jackson ImmunoResearch Inc., West Grove, PA, USA), and 0.5% Triton X-100 (T8787, Sigma-Aldrich, Burlington, MA, USA) for 1 h at room temperature (RT). Sections were then washed twice in PBS for 5 min each, and incubated with Fab fragments (115-007-003, Jackson ImmunoResearch Inc., West Grove, PA, USA) for 30 min to reduce non-specific binding of secondary antibodies, followed by two additional washes in PBS for 5 min each. Subsequently, sections were incubated with primary antibodies diluted in blocking solution as follows: MyHC I (2/3 dilution, BA-D5, DSHB, Iowa, IA, USA), MyHC IIA (1/3 dilution, SC-71, DSHB, Iowa, IA, USA), MyHC IIB (2/3 dilution, BF-F3, DSHB, Iowa, IA, USA), MyHC IIX (2/3 dilution, 6H1, DSHB, Iowa, IA, USA), Laminin (1/300 dilution, L9393, Sigma-Aldrich, Burlington, MA, USA), Neurofilament NfL C-terminus (1/2,000 dilution, RPCA-NfL-ct, EnCor Biotechnology Inc., Gainesville, FL, USA), neurofilament NfL DegenoTag Peptide (1/2.000 dilution, MCA-6H63, EnCor Biotechnology Inc., Gainesville, FL, USA) or NeuroTrace 500/525 Fluorescent Nissl (1/500, N21480, Invitrogen, Waltham, MA, USA). RPCA-NfL-Ct (healthy NfL antibody) binds to the C-terminal tail of NfL, a region that is destroyed or removed during degeneration thereby labeling healthy neurons; and CPCA-NfL-Degen (degenerated NfL antibody) binds to another region that becomes exposed only in degenerated forms, thus marking damaged neurons. After primary antibody incubation, sections were rinsed in PBS and incubated with the following fluorescent labelled secondary antibodies for 1 h at RT: Alexa Fluor 488 (1/300 dilution, A21042, Invitrogen, Waltham, MA, USA), Alexa Fluor 647 (1/300 dilution, A21240, Invitrogen, Waltham, MA, USA), Cy3 (1/300 dilution, 115-167-186, Jackson ImmunoResearch Labs Inc., Philadelphia, PA, USA), and Cy5 (1/100 dilution, 111-475-003, Jackson ImmunoResearch Labs Inc., Philadelphia, PA, USA). Sections were counterstained with 4′,6-diamidino-2-phenylindole (DAPI). Finally, slides were mounted using Fluoromount-G.

### Optical imaging and analysis

TA tissue histology and immunofluorescence staining were scanned using a Zeiss Axioscan 7 microscope (Zeiss, Oberkochen, Germany), while spinal cord samples were imaged using a Zeiss LSM900 confocal microscope (Zeiss, Oberkochen, Germany). Myofiber analysis was performed using the MuscleJ2 plugin ^71^ in Fiji (ImageJ v2.18.0/1.54p) ^72^. Cross-sectional area (CSA) was measured and represented either as an average or as a histogram distribution. Fiber type identification was performed based on MYH expression. Spinal cord images were analyzed in Fiji, where the integrated density of different NfL epitopes was quantified in damaged and healthy neurons, and the ratio between the corresponding signals was calculated.

### LIMS experiments

Fresh-frozen 14-µm-thick cryosections were mounted on indium-tin oxide (ITO)-coated slides (Intellislides, Bruker Daltonics, Germany), and stored at -80 °C. After thawing, slides were covered with 1,5-diaminonaphthalene (DAN) using an in-house designed sublimator ^49^. Sections were scanned in negative polarity, using the timsTOF fleX mass spectrometer (MS; Bruker Daltonics, Germany) available at the analytical core facility of the University of the Basque Country (UPV/EHU, Spain). Negative polarity was used to detect a larger number of lipid classes. Indeed, we identified phosphatidylinositols (PI), phosphatidylethanolamines (PE), phosphatidylglycerols (PG), phosphatidylserines (PS), sphingomyelins (SM) and sulfatides (SFT). In addition to their ether and lyso variants: ether-linked lyso-phosphatidylethanolamine (LPEe), ether-linked lyso-phosphatidylcholine (LPCe), LPIe, ether-linked lysophosphatidylinositols (LPIe), LPE, lyso-phosphatidylethanolamines (LPE), lyso-phosphatidylcholines (LPC), lyso-phosphatidylglycerols (LPG), lyso-phosphatidylinositols (LPI), lysosphingomyelins (LSM), and lysosulfatides (LSFT). Due to the existence of multiple isobaric species, PC/PE are presented together and PCe/PEe grouped in a single graph.

Data acquisition was performed at 10 µm/pixel using 100 shots/pixel with a laser energy of approximately 40 μJ/pulse. A mass observation window of 350-1,300 Da was used, within which the MS achieves a mass resolution of approximately 60,000 at m/z = 1000. Spectra were processed using either previously described software developed in MATLAB (MathWorks, R2026a, Nantick, MA, USA) ^49,73^ or SCiLS (Bruker Daltonics, version 2025b). Concisely, our in-house software aligns the spectra, applies total ion current normalization, and calculates an average spectrum, which is then used to extract the peaks with intensity above 0.5% of the strongest peak ^73^. The resulting peak list was processed by removing unwanted adducts, and only those m/z values corresponding to the lipids of interest were used, which significantly reduced the size of the dataset without a significant loss of information and prepared the data for further analysis. Images were then segmented using the lipid fingerprint at each pixel and an unsupervised divisive hierarchical clustering-rank compete (DHC-RC) algorithm. In parallel, SCiLS applied supervised k-means for image segmentation. After obtaining the complete list of peaks, each pairwise comparison was analyzed independently. First, a lipid family analysis was performed by summing the abundances of all lipid species belonging to the same family to obtain the total abundance per family. Subsequently, a principal component analysis (PCA) was conducted in RStudio (Posit Software, Boston, MA, USA; version 2024.12.0+467) to visualize the statistically significant lipid species across comparisons.

### HPLC analysis

High-performance liquid chromatography (HPLC) analysis was performed at the Mass Spectrometry and Molecular Imaging Unit (IMSMI), Maimonides Biomedical Research Institute of Córdoba (IMIBIC, Spain). Serum from male (n=5) and female (n=3) WT and mutant mice was collected by cardiac puncture into BD Microtainer tubes with separating gel (365968, BD Biosciences, Franklin Lakes, NJ, USA) and kept at 4 °C. Tubes were then centrifuged at 6,000 x g for 10 min at 4 °C. After centrifugation, supernatant was collected, frozen in liquid nitrogen, and stored at -80 °C until further use.

Serum samples were thawed on ice prior to processing. For each sample, 50 μL of serum was mixed with 300 μL of isopropanol at RT, incubated for 15 min at RT, and then centrifuged at 12,000 × g for 10 min at 10 °C. After centrifugation, supernatant was collected, and an additional centrifugation step was performed if necessary to ensure homogeneity.

Liquid chromatography (LC) separation was performed using an Elute UHPLC system (Bruker Daltonics, Germany) with mobile phase A (ACN:H₂O, 60:40) and mobile phase B (IPA:ACN:H₂O, 85:10:5), both containing 0.1% formic acid (Thermo Fisher Scientific, Waltham, MA, USA) and 10 mM ammonium formate (Merck, Darmstadt, Germany). The LC gradient consisted of a 20-min runtime as follows: T0: 60% A/40% B; T0-T2: 57% A/43% B; T2-T2.10: 50% A/50% B; T2.10-T12: 46% A/54% B; T12-T12.10: 30% A/70% B; T12.10-T15: 1% A/99% B; T15-T18.10: 60% A/40% B; T18.10-T20: 60% A/40% B. Samples were injected in triplicate using the μL pickup mode with a 5 μL injection volume. Separation was performed on an Acquity PRM BEH C18 1.7 μm, 2.1 × 100 mm column coupled to an AQTY PRM BEH C18 1.7 μm, 2.1 × 5 mm pre-column (Waters Cromatografía, S.A., Barcelona, Spain), thermostated at 55 °C.

4D PASEF (parallel accumulation serial fragmentation) experiments were conducted using a TIMS-TOF Pro instrument coupled to a VIP-HESI source (Bruker Daltonics, Germany) in positive and negative polarity. Instrument parameters were as follows: end plate offset 500 V; capillary voltage, 4,500 V; dry gas (N₂) at 8 L/min at 230 °C; nebulizer gas 2 bar; peak detection threshold 100 counts; PASEF scan mode, mass range, 100-1,350 Da for MS and MS²; acquisition cycle 0.1 s, mobility range 0.55–1.90 Vs/cm²; collision energy 10 eV. TIMS and mass calibration were performed weekly using the Agilent ESI LC-MS tuning mix. Online recalibration was performed immediately after each sample using a mixture of Agilent ESI LC-MS tune mix and 1 mM sodium formate (1:1), injected directly into the ESI source via a syringe pump.

Initial data evaluation was carried out using DataAnalysis 6.1 (Bruker Daltonics). Untargeted data processing was performed in Bruker Compass MetaboScape 2026 v16.0.6 (Bruker Daltonics) using the T-ReX algorithm for mass calibration, peak picking, time alignment, and within-batch correction. All samples were analyzed in triplicate with randomized injection order. Pooled quality control (QC) samples, prepared as an equimolar pool of all study samples, were injected every 9 samples to enable signal normalization and drift correction. Missing values were addressed using the T-ReX automatic region complete extraction algorithm, which re-extracts features below the detection threshold in a targeted manner. Only features detected in all QC samples were retained for further analysis.

Serial dilutions of the QC sample (20, 40, 60, 80, and 100% of the original concentration) were used to evaluate the linearity of the instrument response for each detected feature, using a custom quality-control R script developed by the IMSMI staff. For each feature, all possible aggregation strategies for replicate injections per dilution level were evaluated (first, second, third injection, and the mean), with and without excluding the lowest dilution point (drop-lowest strategy). The combination yielding the highest coefficient of determination (R²) among those with a positive slope was selected. Only features with a positive slope and R² ≥ 0.75 in positive and negative mode were retained, ensuring that only analytically reliable signals were considered for downstream analysis. For features with exactly two valid dilution points, a simplified criterion was applied: acceptance was granted if the slope was positive, without requiring a minimum R². Features with a negative slope but R² ≥ threshold were flagged as potential ion suppression candidates and retained with an “ion suppression” label. Additionally, a repeatability filter was applied: only features with a coefficient of variation (%CV) across injection replicates ≤ 20% were retained. Injection replicates were first screened for global failure (total intensity sum and median intensity below 70% of the replicate median), and metabolite-level outliers (|Z-score| > 2) were excluded before averaging.

### RNA extraction and RNA-seq analysis

Total RNA was isolated from quadriceps muscle tissue of control and *hSOD1^G93A^* mice at > P118. Tissue dissociation was performed using a bead homogenizer (19-050A, Omni, Kennesaw, GA, USA), and RNA extraction was then carried out using the Maxwell RSC simplyRNA Tissue Kit (AS1340, Promega, Madison, WI, USA) according to the manufacturer’s instructions.

RNA concentration was measured via fluorometric quantification using a Qubit 4 instrument (Thermo Fisher Scientific), and RNA integrity was assessed with the Agilent RNA 6000 Nano Kit on a 2100 Bioanalyzer (Agilent Technologies). All samples exhibited high RNA quality, with RNA integrity numbers (RIN) above 7. mRNA molecules were selectively captured and purified using the Dynabeads mRNA Purification Kit (Thermo Fisher Scientific). Strand-specific RNA libraries were prepared using the MGIEasy Fast RNA Library Prep Set (MGI) on a MGISP100 platform, and library quality was evaluated with the Agilent High Sensitivity DNA Kit on the 2100 Bioanalyzer. Next-generation sequencing was performed on a DNBSEQ-G400 platform (paired-end, 150 bp reads, 40 million reads per sample).

RNA-seq data processing and differential expression analysis were performed on the Genomic Platform of the Biogipuzkoa HRI using the standardized nf-core/rnaseq (v3.15.0) and nf-core/differential abundance (v1.5.0) pipelines to ensure reproducibility and best practices in computational workflows ^74^. Raw sequencing data were processed through nf-core/rnaseq. Low-quality base filtering (Phred score < 15) using a sliding window approach and adapter removal were performed using fastp v.0.23.4 ^75^. Ribosomal RNA (rRNA) was removed using SortMeRNA v.4.3.6 ^76^. Cleaned reads were aligned to the *Mus musculus* GRCm39 reference genome (ENSEMBL) using STAR aligner v.2.7.10a ^77^. The full sequence for the *Homo sapiens* SOD1 gene (ENSEMBL ID: ENSG00000142168) was concatenated to the mouse reference genome index prior to alignment to enable accurate quantification of the human transgene. Transcript abundance was quantified at the gene level with Salmon v.1.10.1 ^78^. Count matrices generated by nf-core/rnaseq were analyzed for differential expression using nf-core/differential abundance, which leverages DESeq2 (v1.44.0) ^79^. *p*-values were adjusted using the Benjamini-Hochberg method to reduce false positives. Differentially expressed genes were called with thresholds of log2(fold change) ≥ 1 and adjusted *p*-value (false discovery rate, FDR) < 0.05. Over-representation analysis (ORA) was performed using the enrichGO function from the R package clusterProfiler (v.4.12.6) ^80^. Only significant enrichment results (adjusted *p*-value ≤ 0.05) were considered.

### Statistical analysis

Normality was first assessed using the Shapiro-Wilk test, and homogeneity of variances was verified with Levene’s test. Based on these results, for normally distributed data with equal variances, one-way ANOVA followed by Tukey’s HSD post hoc test was applied. When variances were unequal, Welch ANOVA followed by Games-Howell post hoc test was used. For non-normally distributed data, the Kruskal-Wallis test followed by Wilcoxon rank-sum test was applied. All data are presented as mean ± standard error of the mean (SEM). Statistical significance is indicated using standard asterisk notation: \**P* < 0.05, \*\*\**P* < 0.001, and \*\**P* < 0.01.

Four supervised classification models, namely least absolute shrinkage and selection operator (LASSO), support vector machine (SVM), linear discriminant analysis (LDA), and random forest (RF), were implemented using Orange software (version 3.40). The selected lipid biomarkers were used as input features for sample classification, and the classification performance obtained with each algorithm was subsequently compared. The classification results are presented in terms of the area under the receiver operating characteristic curve (AUC), sensitivity (SEN), and specificity (SPE).

## Data availability

RNA-seq data reported in this publication have been deposited in NCBI’s Gene Expression Omnibus (GEO) and are accessible through GEO series accession number GSE330670.

## Acknowledgements

The authors acknowledge the technical and human support of the Histology, Genomics and Animal Facility Platforms of Biogipuzkoa Health Research Institute, the mass spectrometry resources of SGIker (EHU), the IMIBIC Mass Spectrometry and Molecular Imaging Unit of the Maimonides Biomedical Research Institute of Córdoba, and the Animal Facility of Centro de Investigación Biomédica de Aragón.

## Author information

This project was administered by SA-M. SA-M conceived, planned and supervised these experiments. MG-P, AV-G, OP-M, ML, AE, IA, LR-G, conducted laboratory experiments. MG-P and CH performed LIMS experiments. MG-P, CH, AV-G, OP-M, performed data analysis. LM-M and RO provided mice for the experiments. MR-H performed RNA-seq analysis and interpretation. MG-P, AE, ALM, JAF and SA-M contributed to funding acquisition and supported the project. MG-P and SA-M wrote, reviewed and edited the original draft. JAF revised critically the original draft and provide lipid interpretation of the results. All authors have read, reviewed and approved the final version of this paper.

