## Supplemental_Figures for "Muscle spatial lipidomics identifies early ALS signatures in presymptomatic SOD1G93A mice"

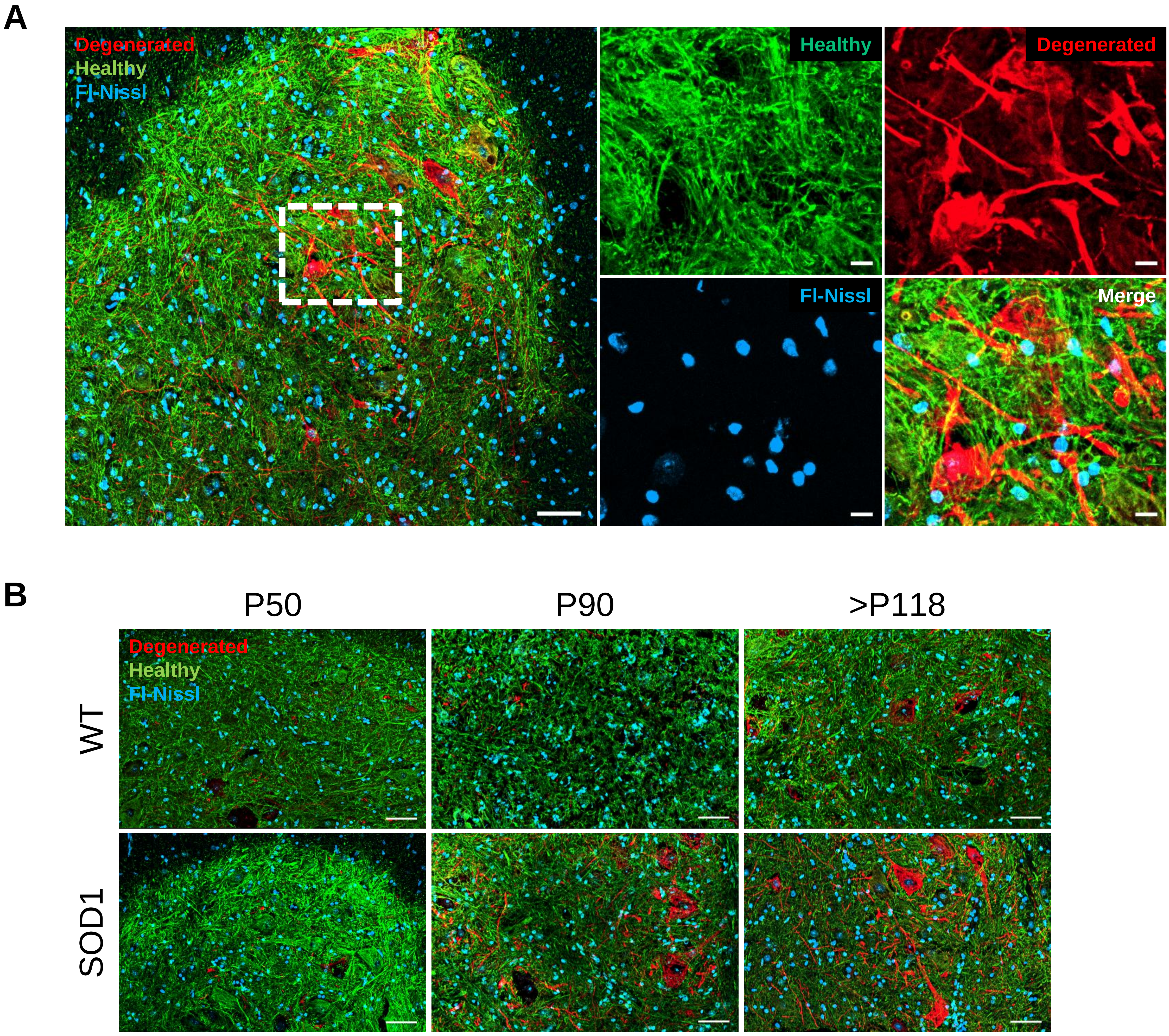


**Supplemental Figure 1.** (**A**) Representative immunofluorescence images of a ventral horn of the spinal cord from end-stage female *hSOD1$^{G93A}$* mutant mice stained in green with RPCA-NF-L-ct (marker of **healthy** NF-L isoform), in red with CPCA-NF-L-Degen (marker of **degenerated** NF-L isoform), and in blue (nuclei) with DAPI Scale bar = 50 µm. White-dot square is magnified on the right. Scale bar = 10 µm. (**B**) Representative immunofluorescence of a ventral horn of a spinal cord from WT and *hSOD1$^{G93A}$* female mutant mice at different disease stages in blue (neurons nuclei) with Fluoro-Nissl. Scale bars = 50 µm. P50, P90 or >P118, correspond to postnatal days 50 (presymptomatic), 90 (symptomatic) or >P118 (end-stage), respectively.

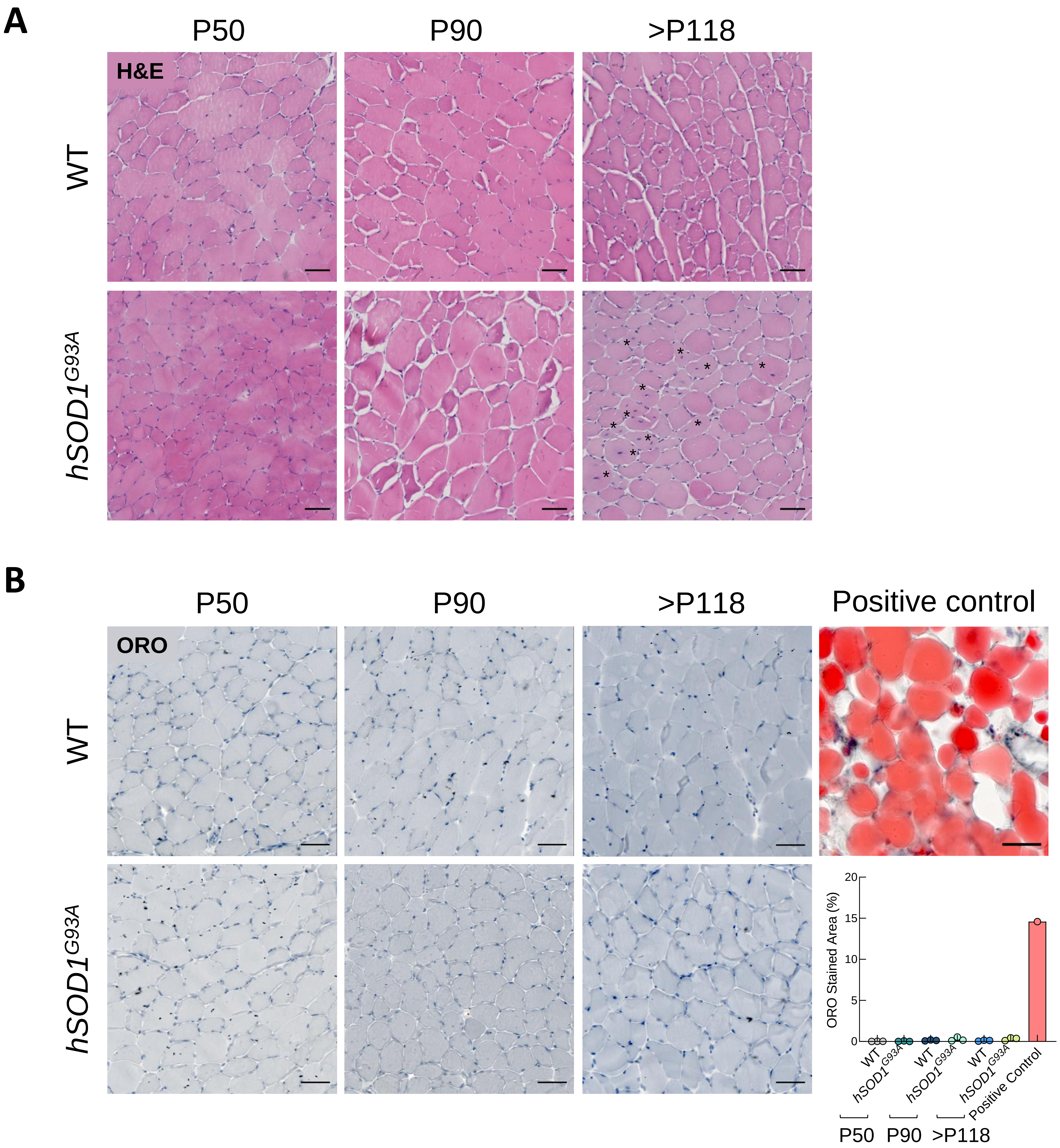


**Supplemental Figure 2.** Representative images of (**A**) Hematoxylin and Eosin (H&E), and (**B**) Oil Red O (ORO) and quantification of the *Tibialis anterior* muscle from wild-type (WT) and *hSOD1*$^{G93A}$ female mice at different disease stages. Black asterisks in H&E indicate myofibers with central-located nuclei. Mouse skin tissue was used as a positive control for ORO staining. Scale bars = 50 µm. P50, P90 or >P118, correspond to postnatal days 50 (presymptomatic), 90 (symptomatic) or >P118 (end-stage), respectively. Bar graphs represent mean ± SEM. N≥3 per condition. For normally distributed data with equal variances, one-way ANOVA followed by Tukey's HSD post hoc test was applied. When variances were unequal, Welch ANOVA followed by Games-Howell post hoc test was used. For non-normally distributed data, the Kruskal-Wallis test followed Wilcoxon rank-sum test was applied. Significant differences are indicated as *$p$<0.05, **$p$<0.01, ***$p$<0.001.

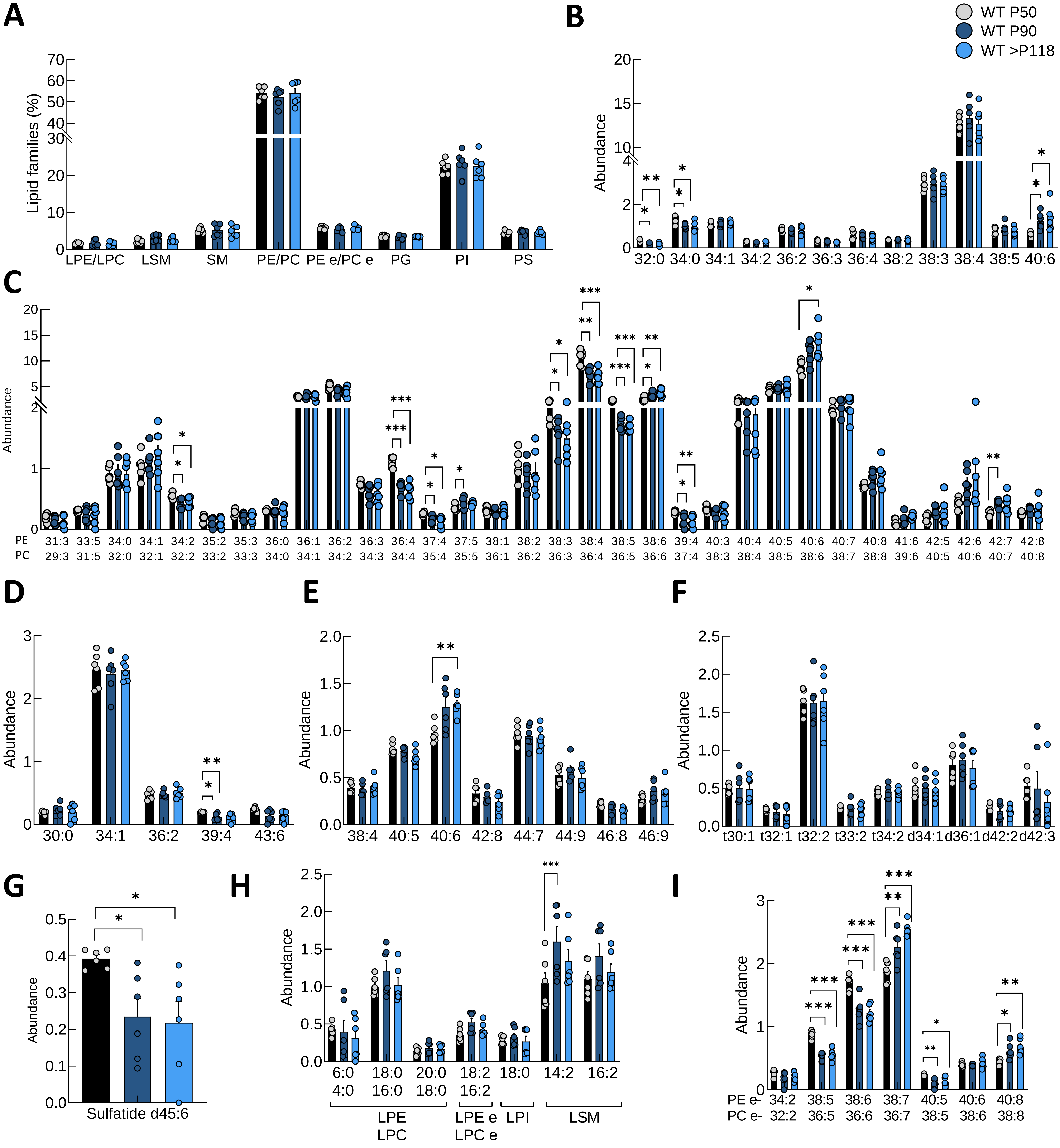


**Supplemental Figure 3.** Relative abundance of (**A**) different lipid families detected, and different lipid species of (**B**) PI, (**C**) PE/PC, (**D**) PG, (**E**) PS, (**F**) SM, (**G**) SFT, (**H**) LPE, LPC, LPEe, LPCe, LPI and LSM, and (**I**) PEe and PCe in *Tibialis anterior* muscle from wild-type female mice at postnatal days 50 (presymptomatic), 90 (symptomatic) or >P118 (end-stage). Abbreviations: PI, phosphatidylinositols; PE, phosphatidylethanolamines; PC, phosphatidylcholines; PG, phosphatidylglycerols; PS, phosphatidylserines; SM, sphingomyelins; SFT, sulfatides; LPE, lysophosphatidylethanolamines; LPC, lysophosphatidylcholines; LPEe, ether-linked lysophosphatidylethanolamines; LPCe, ether-linked lysophosphatidylcholines; LPI, lysophosphatidylinositols; LSM, lysosphingomyelins; PEe, ether-linked phosphatidylethanolamines; PCe, ether-linked phosphatidylcholines. Bar graphs represent mean ± SEM. N≥5 per condition. For normally distributed data with equal variances, one-way ANOVA followed by Tukey's HSD post hoc test was applied. When variances were unequal, Welch ANOVA followed by Games-Howell post hoc test was used. For non-normally distributed data, the Kruskal-Wallis test followed Wilcoxon rank-sum test was applied. Significant differences are indicated as *$p$<0.05, **$p$<0.01, ***$p$<0.001.

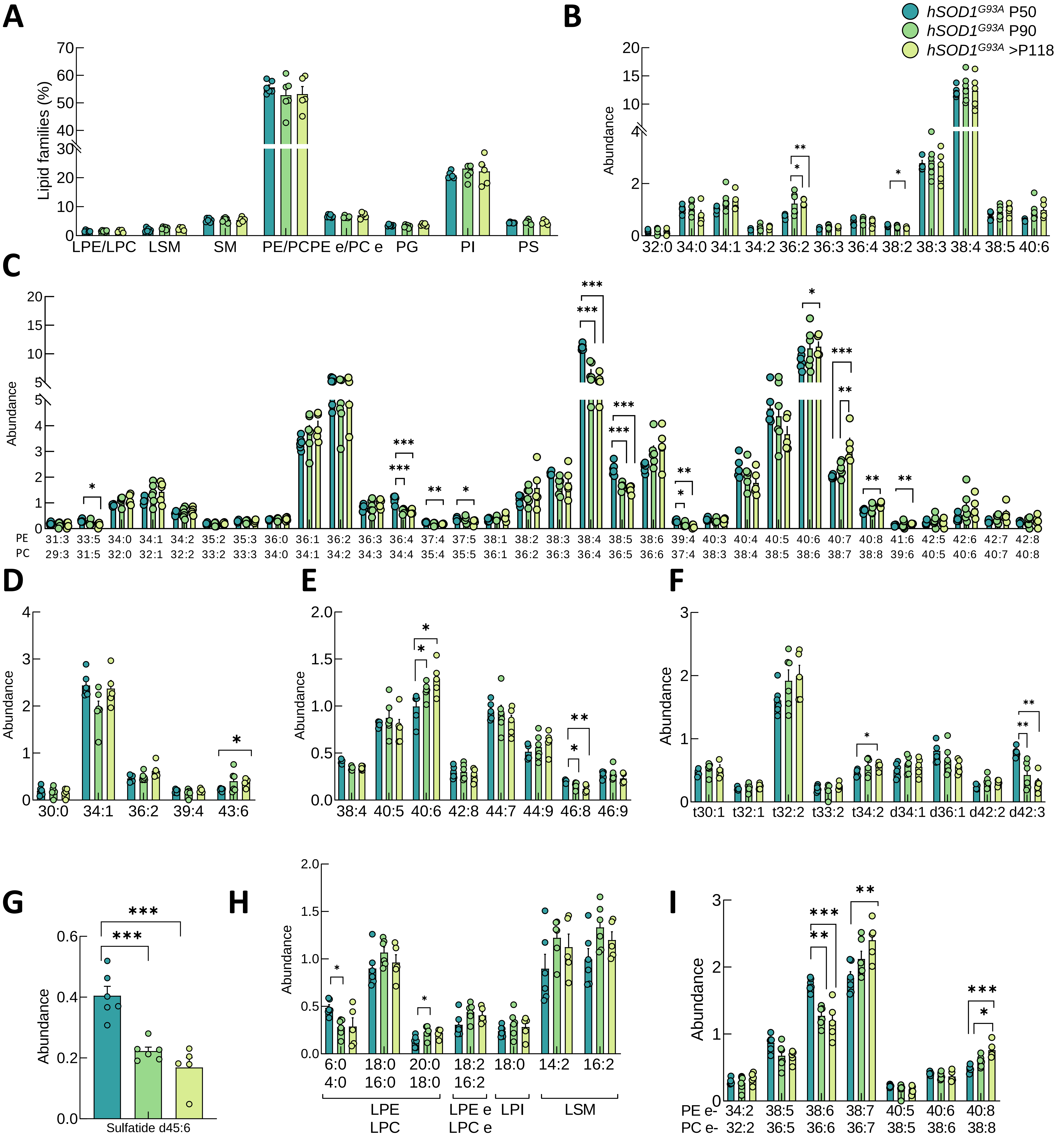


**Supplemental Figure 4.** Relative abundance of (**A**) different lipid families detected, and different lipid species of (**B**) PI, (**C**) PE/PC, (**D**) PG, (**E**) PS, (**F**) SM, (**G**) SFT, (**H**) LPE, LPC, LPEe, LPCe, LPI and LSM, and (**I**) PEe and PCe in *Tibialis anterior* muscle from *hSOD1$^{G93A}$* female mice at postnatal days 50 (presymptomatic), 90 (symptomatic) or >P118 (end-stage). Abbreviations: PI, phosphatidylinositols; PE, phosphatidylethanolamines; PC, phosphatidylcholines; PG, phosphatidylglycerols; PS, phosphatidylserines; SM, sphingomyelins; SFT, sulfatides; LPE, lysophosphatidylethanolamines; LPC, lysophosphatidylcholines; LPEe, ether-linked lysophosphatidylethanolamines; LPCe, ether-linked lysophosphatidylcholines; LPI, lysophosphatidylinositols; LSM, lysosphingomyelins; PEe, ether-linked phosphatidylethanolamines; PCe, ether-linked phosphatidylcholines. Bar graphs represent mean ± SEM. N≥5 per condition. For normally distributed data with equal variances, one-way ANOVA followed by Tukey's HSD post hoc test was applied. When variances were unequal, Welch ANOVA followed by Games-Howell post hoc test was used. For non-normally distributed data, the Kruskal-Wallis test followed Wilcoxon rank-sum test was applied. Significant differences are indicated as *$p$<0.05, **$p$<0.01, ***$p$<0.001.

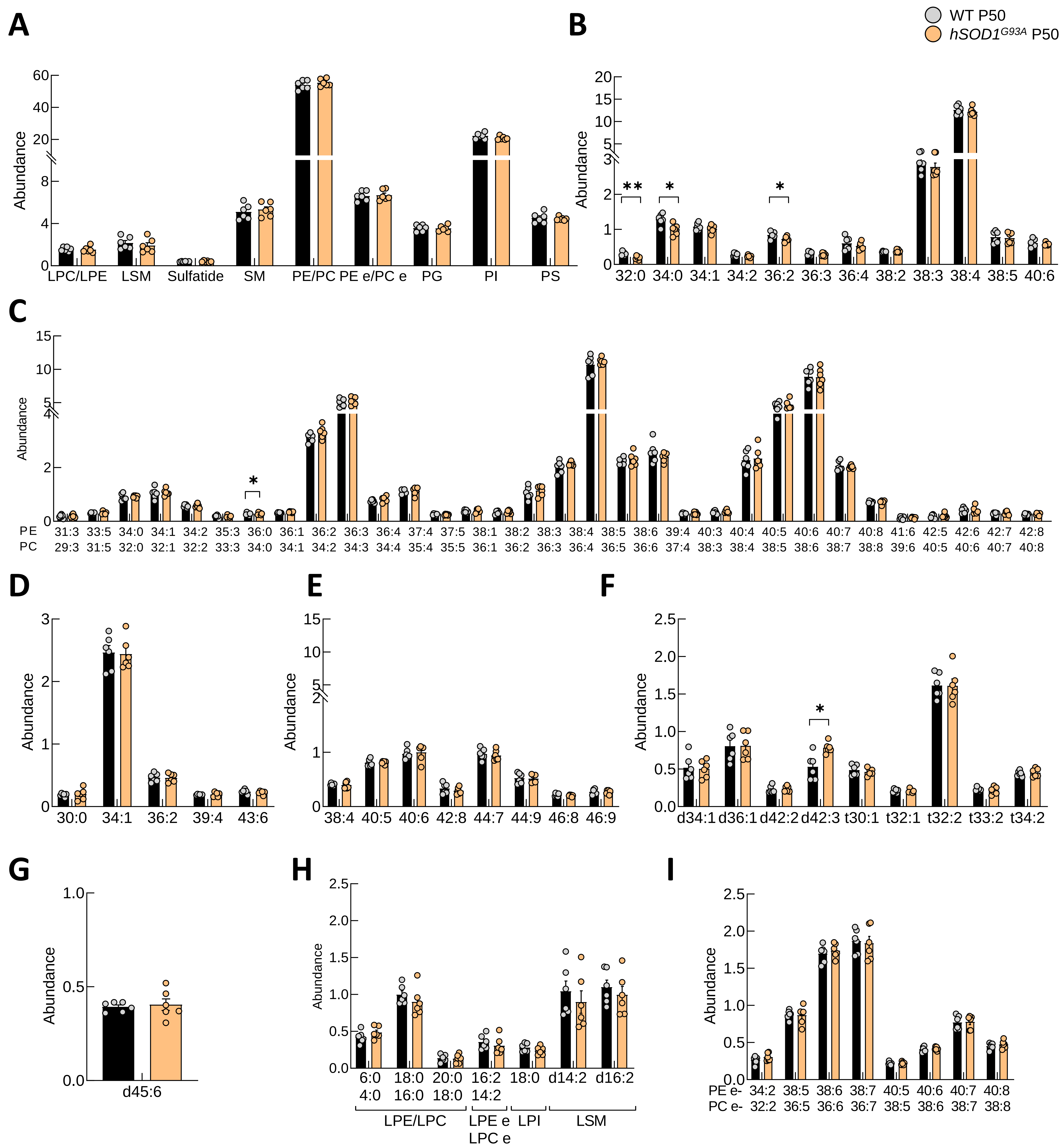


**Supplemental Figure 5.** Relative abundance of (**A**) different lipid families detected, and different lipid species of (**B**) PI, (**C**) PE/PC, (**D**) PG, (**E**) PS, (**F**) SM, (**G**) SFT, (**H**) LPE, LPC, LPEe, LPCe, LPI and LSM, and (**I**) PEe and PCe in *Tibialis anterior* muscle from wild-type and *hSOD1*$^{G93A}$ female mice at postnatal days 50 (presymptomatic), 90 (symptomatic) or >P118 (end-stage). Abbreviations: PI, phosphatidylinositols; PE, phosphatidylethanolamines; PC, phosphatidylcholines; PG, phosphatidylglycerols; PS, phosphatidylserines; SM, sphingomyelins; SFT, sulfatides; LPE, lysophosphatidylethanolamines; LPC, lysophosphatidylcholines; LPEe, ether-linked lysophosphatidylethanolamines; LPCe, ether-linked lysophosphatidylcholines; LPI, lysophosphatidylinositols; LSM, lysosphingomyelins; PEe, ether-linked phosphatidylethanolamines; PCe, ether-linked phosphatidylcholines. Bar graphs represent mean ± SEM. N=6 per condition. For normally distributed data with equal variances, one-way ANOVA followed by Tukey's HSD post hoc test was applied. When variances were unequal, Welch ANOVA followed by Games-Howell post hoc test was used. For non-normally distributed data, the Kruskal-Wallis test followed Wilcoxon rank-sum test was applied. Significant differences are indicated as *$p<0.05$, **$p<0.01$, ***$p<0.001$.

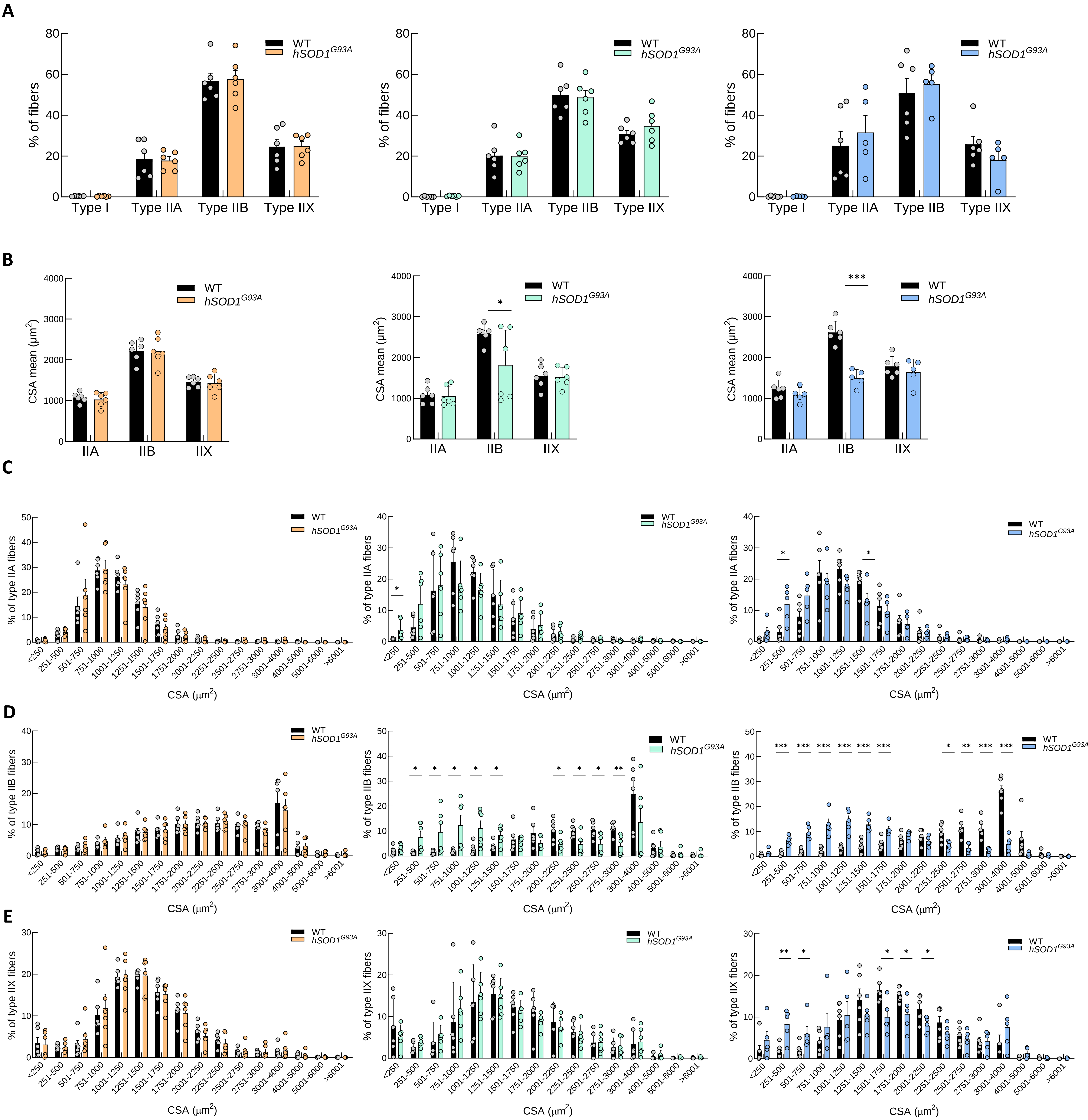


**Supplemental Figure 6.** (**A**) Quantification of type IIA, IIB, and IIX fibers, and (**B**) cross-sectional area (CSA) in *Tibialis anterior* (TA) muscle from wild-type (WT) and *hSOD1$^{G93A}$* female mice across different disease stages. Frequency distribution of CSA for type (**C**) IIA, (**D**) IIB, and (**E**) IIX fibers at postnatal days 50 (presymptomatic), 90 (symptomatic) and > 118 (end-stage). Bar graphs represent mean ± SEM. N≥5 per condition. For normally distributed data with equal variances, one-way ANOVA followed by Tukey's HSD post hoc test was applied. When variances were unequal, Welch ANOVA followed by Games-Howell post hoc test was used. For non-normally distributed data, the Kruskal-Wallis test followed Wilcoxon rank-sum test was applied. Significant differences are indicated as *$p$<0.05, **$p$<0.01, ***$p$<0.001.

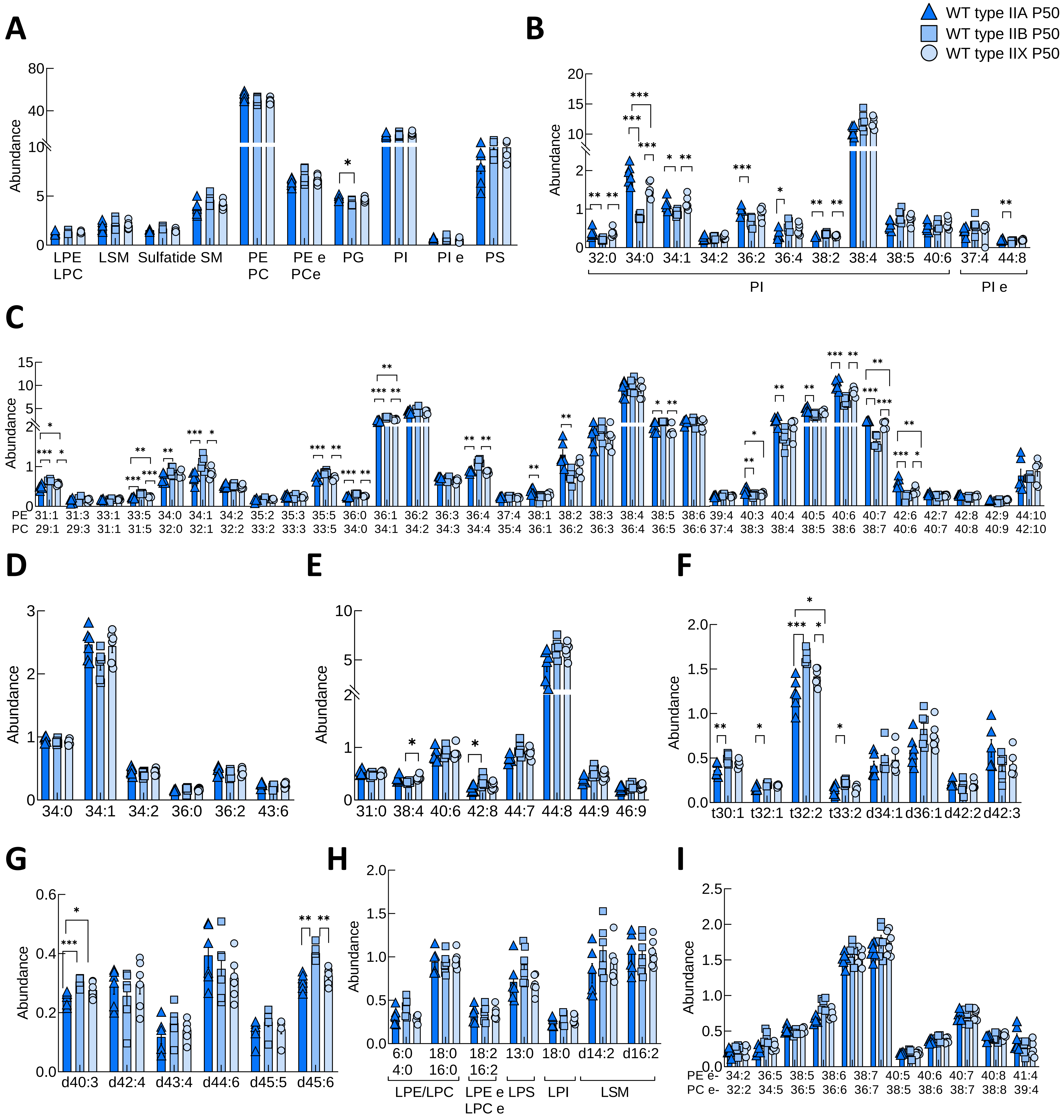


**Supplemental Figure 7.** Relative abundance of (**A**) different lipid families detected, and different lipid species of (**B**) PI and PIe, (**C**) PE/PC, (**D**) PG, (**E**) PS, (**F**) SM, (**G**) SFT, (**H**) LPE, LPC, LPEe, LPCe, LPS, LPI and LSM, and (**I**) PEe and PCe in *Tibialis anterior* muscle from type IIA, IIB, and IIX fibers of wild-type female mice at postnatal days 50 (presymptomatic). Abbreviations: PI, phosphatidylinositols; PIe, ether-linked phosphatidylinositols; PE, phosphatidylethanolamines; PC, phosphatidylcholines; PG, phosphatidylglycerols; PS, phosphatidylserines; SM, sphingomyelins; SFT, sulfatides; LPE, lysophosphatidylethanolamines; LPC, lysophosphatidylcholines; LPEe, ether-linked lysophosphatidylethanolamines; LPCe, ether-linked lysophosphatidylcholines; LPI, lysophosphatidylinositols; LPS, lysophosphatidylserines; LSM, lysosphingomyelins; PEe, ether-linked phosphatidylethanolamines; PCe, ether-linked phosphatidylcholines. Bar graphs represent mean ± SEM. N=6 per condition. For normally distributed data with equal variances, one-way ANOVA followed by Tukey's HSD post hoc test was applied. When variances were unequal, Welch ANOVA followed by Games-Howell post hoc test was used. For non-normally distributed data, the Kruskal-Wallis test followed Wilcoxon rank-sum test was applied. Significant differences are indicated as *$p$<0.05, **$p$<0.01, ***$p$<0.001.

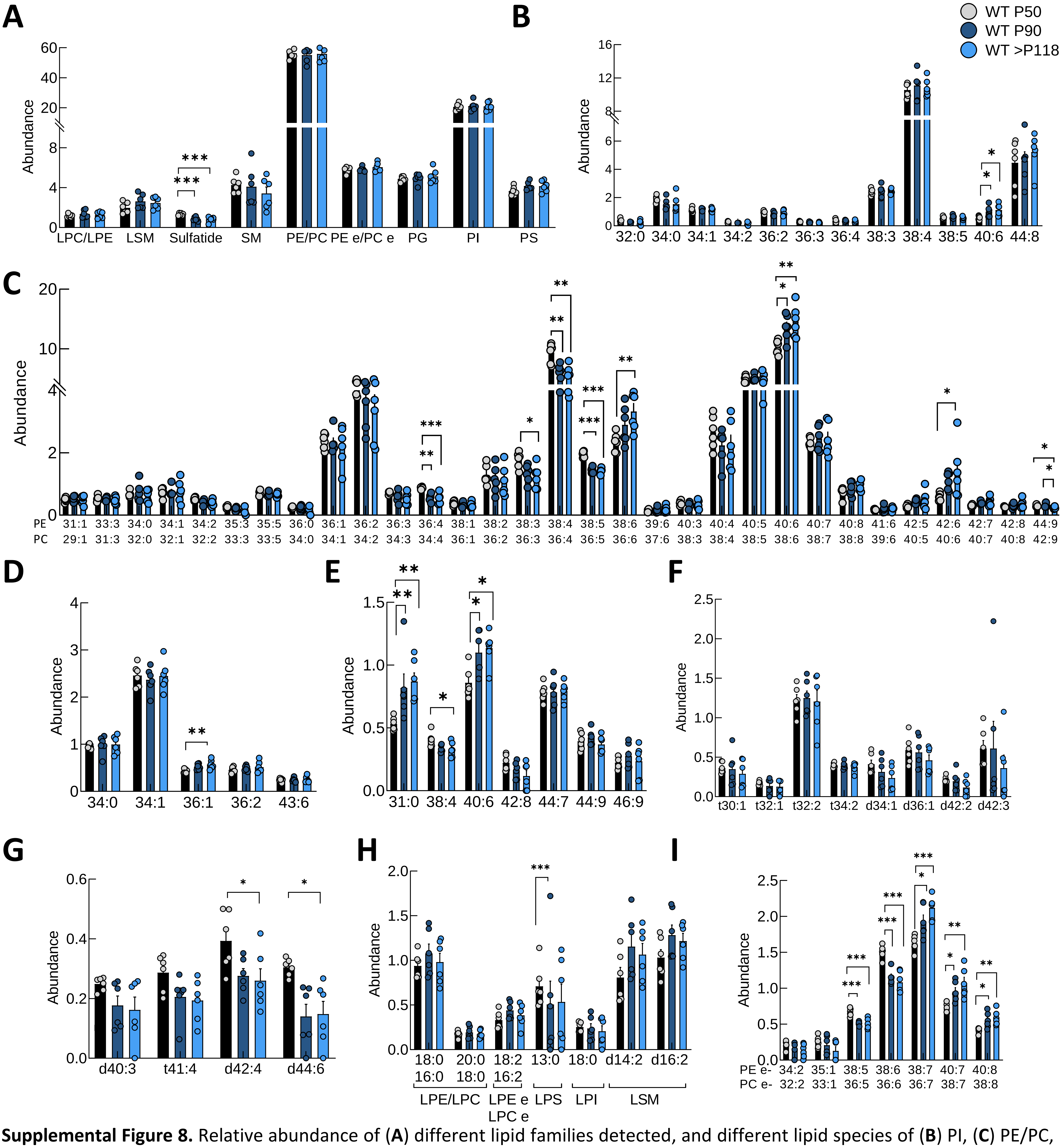


**Supplemental Figure 8.** Relative abundance of (**A**) different lipid families detected, and different lipid species of (**B**) PI, (**C**) PE/PC, (**D**) PG, (**E**) PS, (**F**) SM, (**G**) SFT, (**H**) LPE, LPC, LPEe, LPCe, LPS, LPI and LSM, and (**I**) PEe and PCe in *Tibialis anterior* muscle from IIA fibers of wild-type female mice at postnatal days 50 (presymptomatic), 90 (symptomatic) and > 118 (end-stage). Abbreviations: PI, phosphatidylinositols; PE, phosphatidylethanolamines; PC, phosphatidylcholines; PG, phosphatidylglycerols; PS, phosphatidylserines; SM, sphingomyelins; SFT, sulfatides; LPE, lysophosphatidylethanolamines; LPC, lysophosphatidylcholines; LPEe, ether-linked lysophosphatidylethanolamines; LPCe, ether-linked lysophosphatidylcholines; LPI, lysophosphatidylinositols; LPS, lysophosphatidylserines; LSM, lysosphingomyelins; PEe, ether-linked phosphatidylethanolamines; PCe, ether-linked phosphatidylcholines. Bar graphs represent mean ± SEM. N=6 per condition. For normally distributed data with equal variances, one-way ANOVA followed by Tukey's HSD post hoc test was applied. When variances were unequal, Welch ANOVA followed by Games-Howell post hoc test was used. For non-normally distributed data, the Kruskal-Wallis test followed Wilcoxon rank-sum test was applied. Significant differences are indicated as *$p<0.05$, **$p<0.01$, ***$p<0.001$.

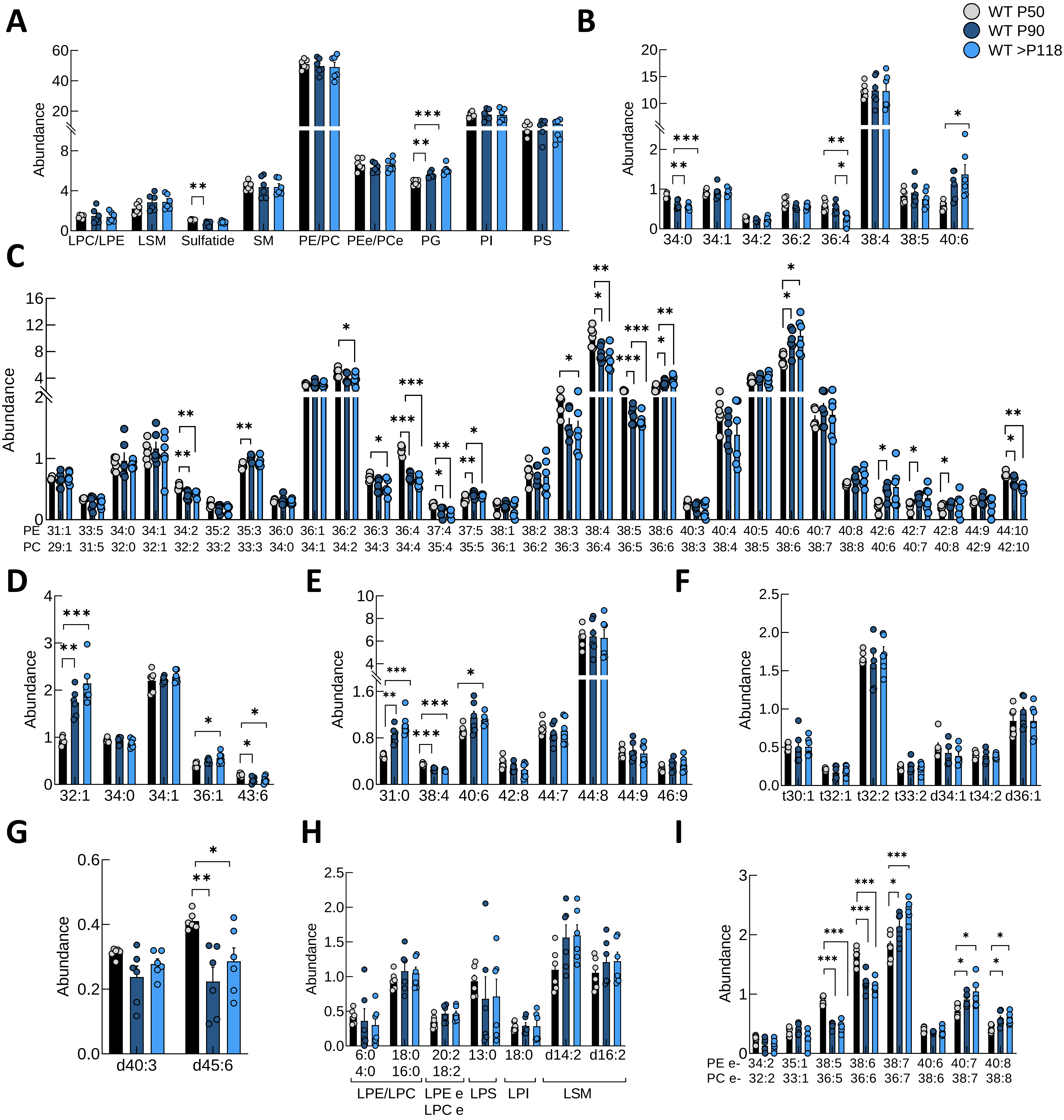


**Supplemental Figure 9.** Relative abundance of (**A**) different lipid families detected, and different lipid species of (**B**) PI, (**C**) PE/PC, (**D**) PG, (**E**) PS, (**F**) SM, (**G**) SFT, (**H**) LPE, LPC, LPEe, LPCe, LPS, LPI and LSM, and (**I**) PEe and PCe in *Tibialis anterior* muscle from IIB fibers of wild-type female mice at postnatal days 50 (presymptomatic), 90 (symptomatic) and > 118 (end-stage). Abbreviations: PI, phosphatidylinositols; PE, phosphatidylethanolamines; PC, phosphatidylcholines; PG, phosphatidylglycerols; PS, phosphatidylserines; SM, sphingomyelins; SFT, sulfatides; LPE, lysophosphatidylethanolamines; LPC, lysophosphatidylcholines; LPEe, ether-linked lysophosphatidylethanolamines; LPCe, ether-linked lysophosphatidylcholines; LPI, lysophosphatidylinositols; LPS, lysophosphatidylserines; LSM, lysosphingomyelins; PEe, ether-linked phosphatidylethanolamines; PCe, ether-linked phosphatidylcholines. Bar graphs represent mean ± SEM. N=6 per condition. For normally distributed data with equal variances, one-way ANOVA followed by Tukey's HSD post hoc test was applied. When variances were unequal, Welch ANOVA followed by Games-Howell post hoc test was used. For non-normally distributed data, the Kruskal-Wallis test followed Wilcoxon rank-sum test was applied. Significant differences are indicated as *$p$<0.05, **$p$<0.01, ***$p$<0.001.

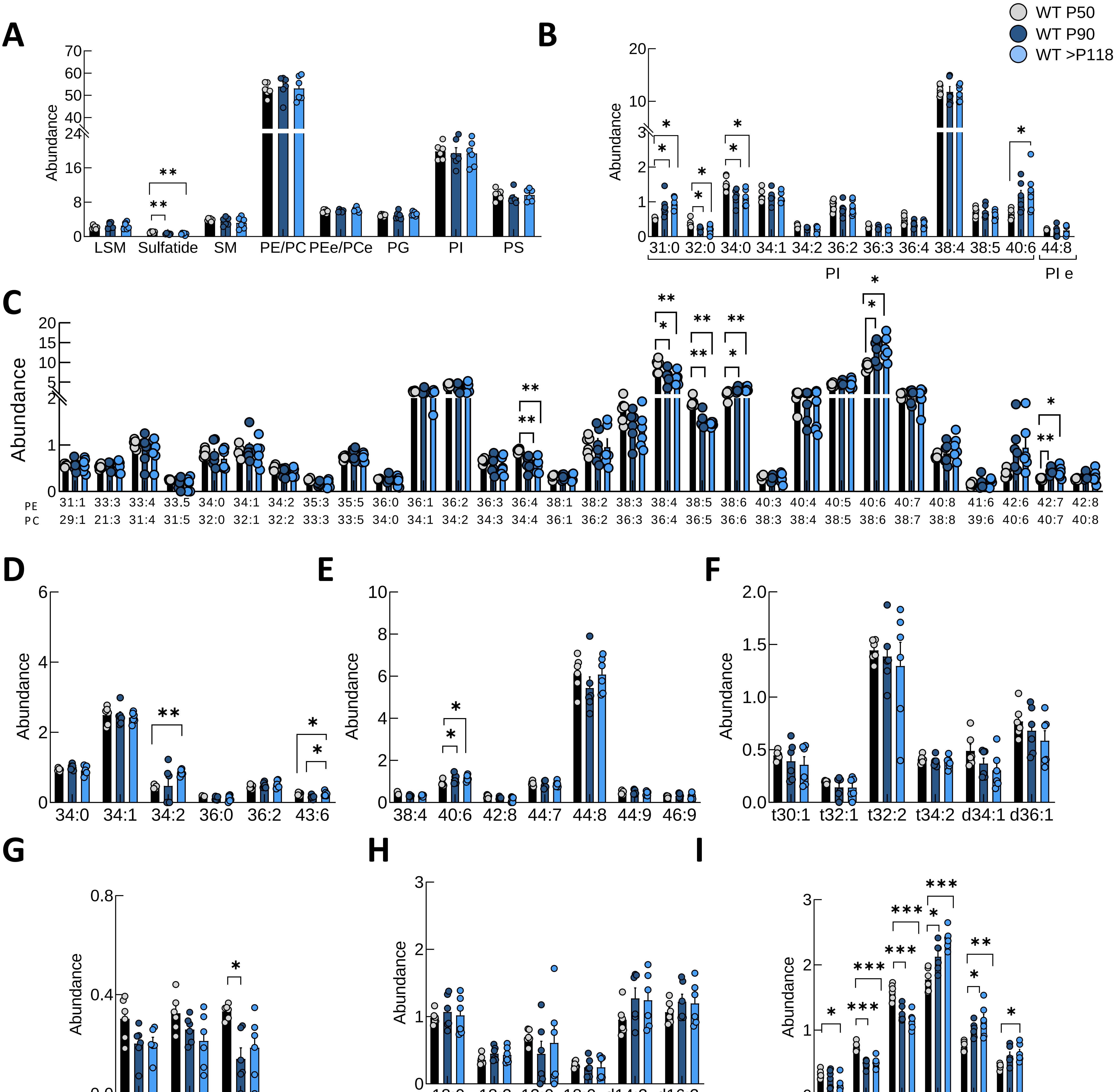


**Supplemental Figure 10.** Relative abundance of (**A**) different lipid families detected, and different lipid species of (**B**) PI and PIe, (**C**) PE/PC, (**D**) PG, (**E**) PS, (**F**) SM, (**G**) SFT, (**H**) LPE, LPC, LPEe, LPCe, LPS, LPI and LSM, and (**I**) PEe and PCe in *Tibialis anterior* muscle from IIX fibers of female wild-type mice at postnatal days 50 (presymptomatic), 90 (symptomatic) and > 118 (end-stage). Abbreviations: PI, phosphatidylinositols; PIe, ether-linked phosphatidylinositols; PE, phosphatidylethanolamines; PC, phosphatidylcholines; PG, phosphatidylglycerols; PS, phosphatidylserines; SM, sphingomyelins; SFT, sulfatides; LPE, lysophosphatidylethanolamines; LPC, lysophosphatidylcholines; LPEe, ether-linked lysophosphatidylethanolamines; LPCe, ether-linked lysophosphatidylcholines; LPI, lysophosphatidylinositols; LPS, lysophosphatidylserines; LSM, lysosphingomyelins; PEe, ether-linked phosphatidylethanolamines; PCe, ether-linked phosphatidylcholines. Bar graphs represent mean ± SEM. N=6 per condition. For normally distributed data with equal variances, one-way ANOVA followed by Tukey's HSD post hoc test was applied. When variances were unequal, Welch ANOVA followed by Games-Howell post hoc test was used. For non-normally distributed data, the Kruskal-Wallis test followed Wilcoxon rank-sum test was applied. Significant differences are indicated as *$p<0.05$, **$p<0.01$, ***$p<0.001$.

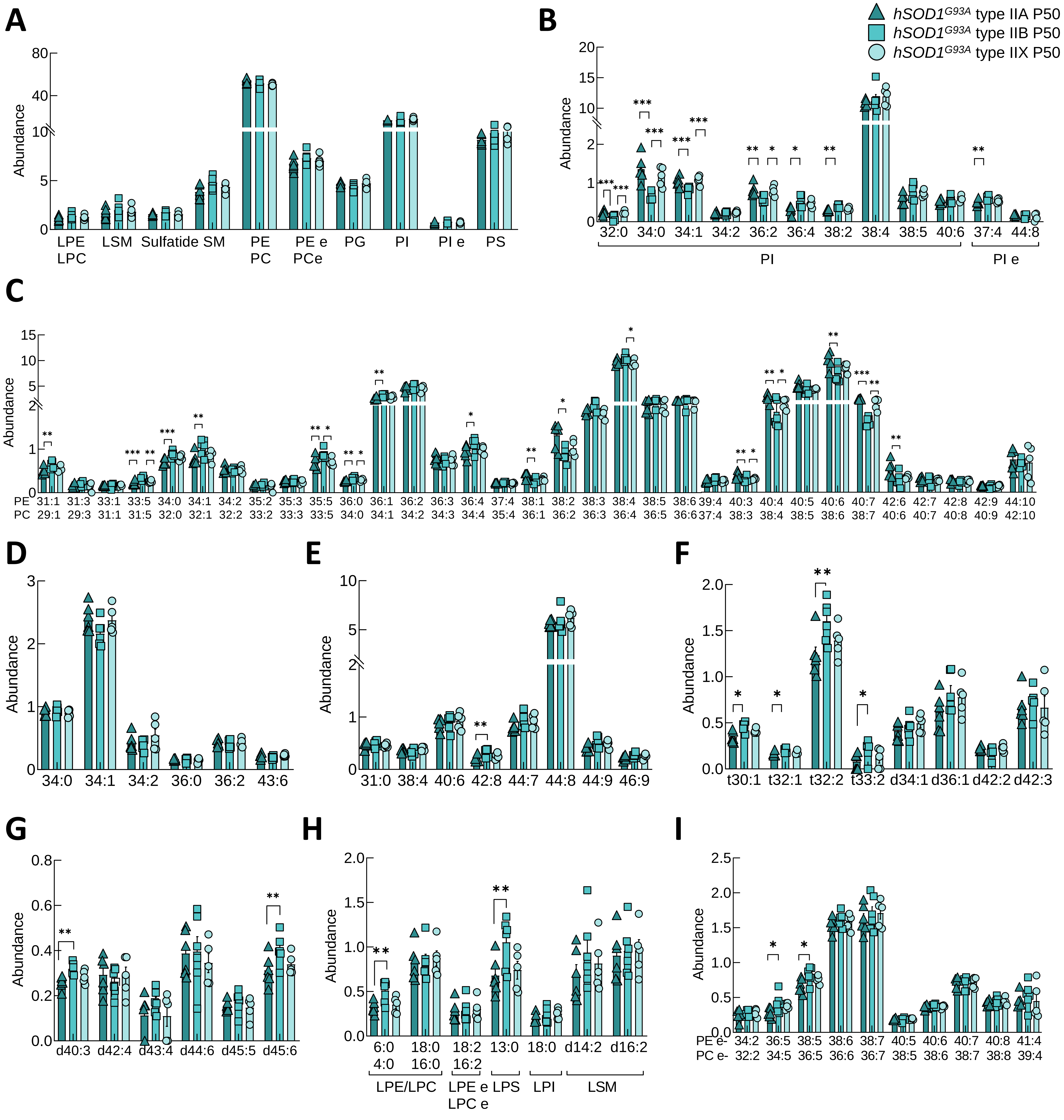


**Supplemental Figure 11.** Relative abundance of (**A**) different lipid families detected, and different lipid species of (**B**) PI and PIe, (**C**) PE/PC, (**D**) PG, (**E**) PS, (**F**) SM, (**G**) SFT, (**H**) LPE, LPC, LPEe, LPCe, LPS, LPI and LSM, and (**I**) PEe and PCe in *Tibialis anterior* muscle from IIA, IIB, and IIX fibers of *hSOD1*$^{G93A}$ female mice at postnatal day 50 (presymptomatic). Abbreviations: PI, phosphatidylinositols; PIe, ether-linked phosphatidylinositols; PE, phosphatidylethanolamines; PC, phosphatidylcholines; PG, phosphatidylglycerols; PS, phosphatidylserines; SM, sphingomyelins; SFT, sulfatides; LPE, lysophosphatidylethanolamines; LPC, lysophosphatidylcholines; LPEe, ether-linked lysophosphatidylethanolamines; LPCe, ether-linked lysophosphatidylcholines; LPI, lysophosphatidylinositols; LPS, lysophosphatidylserines; LSM, lysosphingomyelins; PEe, ether-linked phosphatidylethanolamines; PCe, ether-linked phosphatidylcholines. Bar graphs represent mean ± SEM. N≥5 per condition. For normally distributed data with equal variances, one-way ANOVA followed by Tukey's HSD post hoc test was applied. When variances were unequal, Welch ANOVA followed by Games-Howell post hoc test was used. For non-normally distributed data, the Kruskal-Wallis test followed Wilcoxon rank-sum test was applied. Significant differences are indicated as *$p$<0.05, **$p$<0.01, ***$p$<0.001.

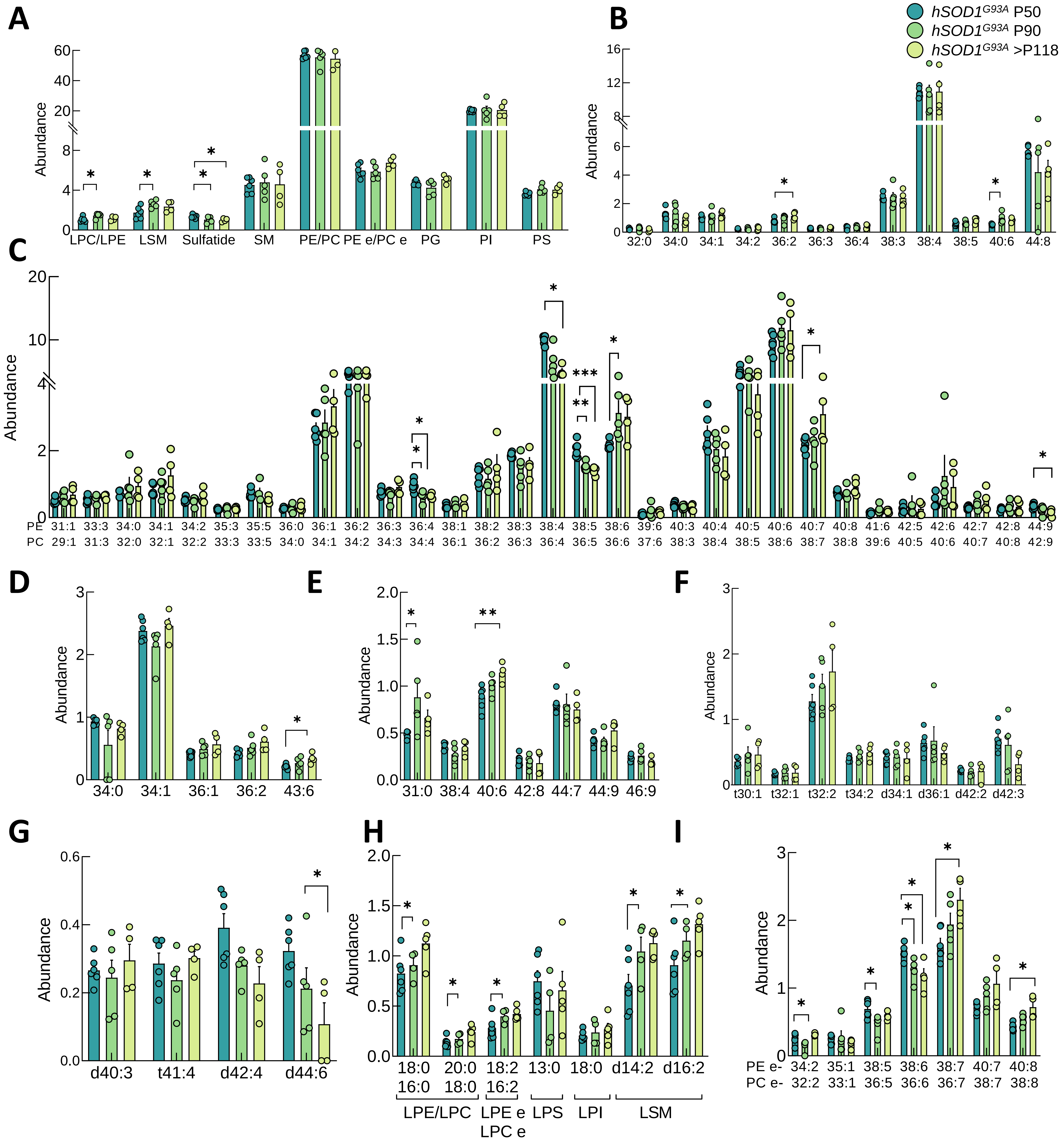


**Supplemental Figure 12.** Relative abundance of (**A**) different lipid families detected, and different lipid species of (**B**) PI, (**C**) PE/PC, (**D**) PG, (**E**) PS, (**F**) SM, (**G**) SFT, (**H**) LPE, LPC, LPEe, LPCe, LPS, LPI and LSM, and (**I**) PEe and PCe in *Tibialis anterior* muscle from IIA fibers of *hSOD1*$^{G93A}$ female mice at postnatal days 50 (presymptomatic), 90 (symptomatic) and > 118 (end-stage). Abbreviations: PI, phosphatidylinositols; PE, phosphatidylethanolamines; PC, phosphatidylcholines; PG, phosphatidylglycerols; PS, phosphatidylserines; SM, sphingomyelins; LPE, lysophosphatidylethanolamines; LPC, lysophosphatidylcholines; LPEe, ether-linked lysophosphatidylethanolamines; LPCe, ether-linked lysophosphatidylcholines; LPI, lysophosphatidylinositols; SFT, sulfatides; LPS, lysophosphatidylserines; LSM, lysosphingomyelins; PEe, ether-linked phosphatidylethanolamines; PCe, ether-linked phosphatidylcholines. Bar graphs represent mean ± SEM. N≥4 per condition. For normally distributed data with equal variances, one-way ANOVA followed by Tukey's HSD post hoc test was applied. When variances were unequal, Welch ANOVA followed by Games-Howell post hoc test was used. For non-normally distributed data, the Kruskal-Wallis test followed Wilcoxon rank-sum test was applied. Significant differences are indicated as *$p<0.05$, **$p<0.01$, ***$p<0.001$.

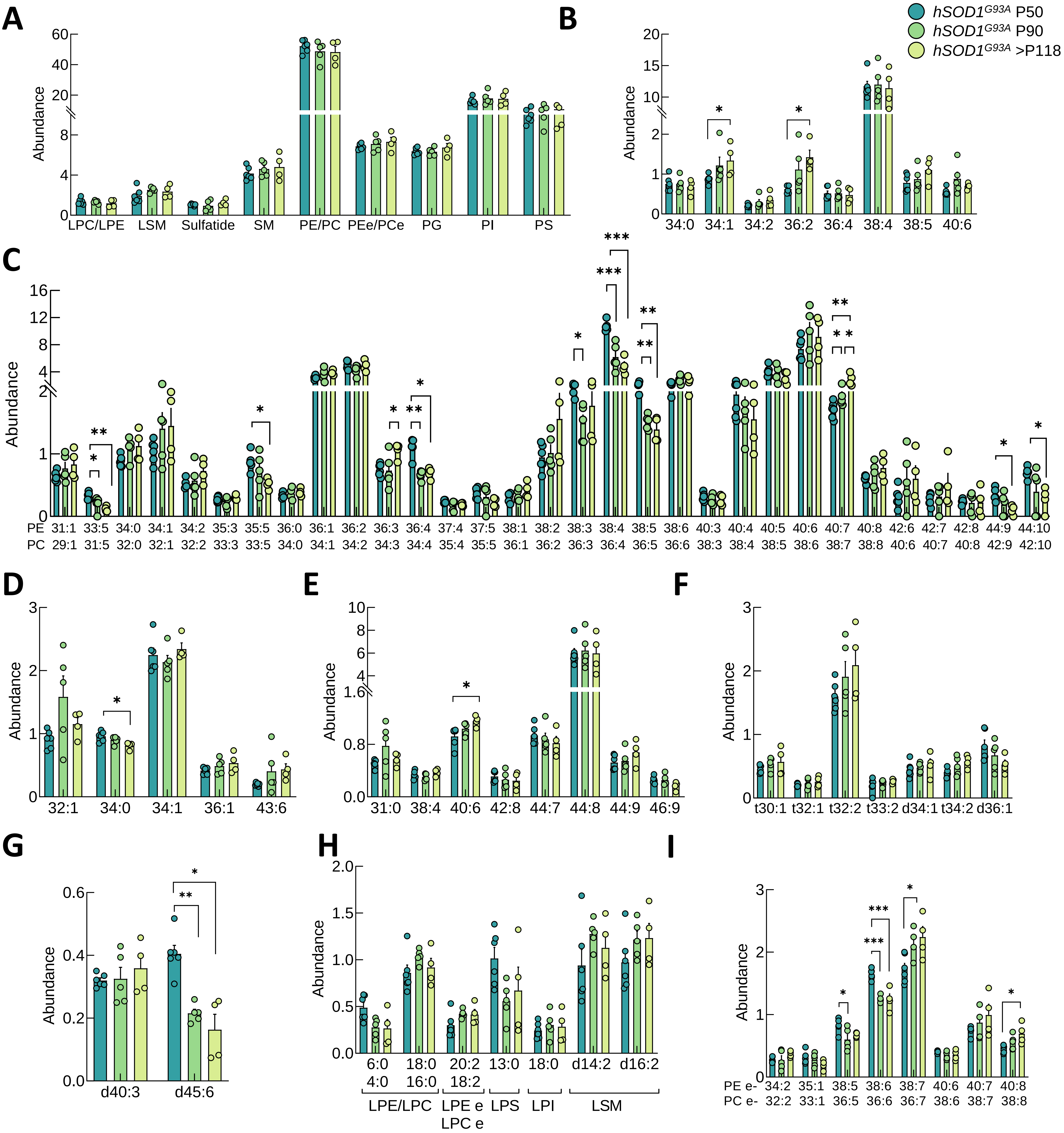


**Supplemental Figure 13.** Relative abundance of (**A**) different lipid families detected, and different lipid species of (**B**) PI, (**C**) PE/PC, (**D**) PG, (**E**) PS, (**F**) SM, (**G**) SFT, (**H**) LPE, LPC, LPEe, LPCe, LPS, LPI and LSM, and (**I**) PEe and PCe in *Tibialis anterior* muscle from IIB fibers of *hSOD1*$^{G93A}$ female mice at postnatal day 50 (presymptomatic), 90 (symptomatic) and > 118 (end-stage). Abbreviations: PI, phosphatidylinositols; PE, phosphatidylethanolamines; PC, phosphatidylcholines; PG, phosphatidylglycerols; PS, phosphatidylserines; SM, sphingomyelins; SFT, sulfatides; LPE, lysophosphatidylethanolamines; LPC, lysophosphatidylcholines; LPEe, ether-linked lysophosphatidylethanolamines; LPCe, ether-linked lysophosphatidylcholines; LPI, lysophosphatidylinositols; LPS, lysophosphatidylserines; LSM, lysosphingomyelins; PEe, ether-linked phosphatidylethanolamines; PCe, ether-linked phosphatidylcholines. Bar graphs represent mean ± SEM. N≥4 per condition. For normally distributed data with equal variances, one-way ANOVA followed by Tukey's HSD post hoc test was applied. When variances were unequal, Welch ANOVA followed by Games-Howell post hoc test was used. For non-normally distributed data, the Kruskal-Wallis test followed Wilcoxon rank-sum test was applied. Significant differences are indicated as *$p$<0.05, **$p$<0.01, ***$p$<0.001.

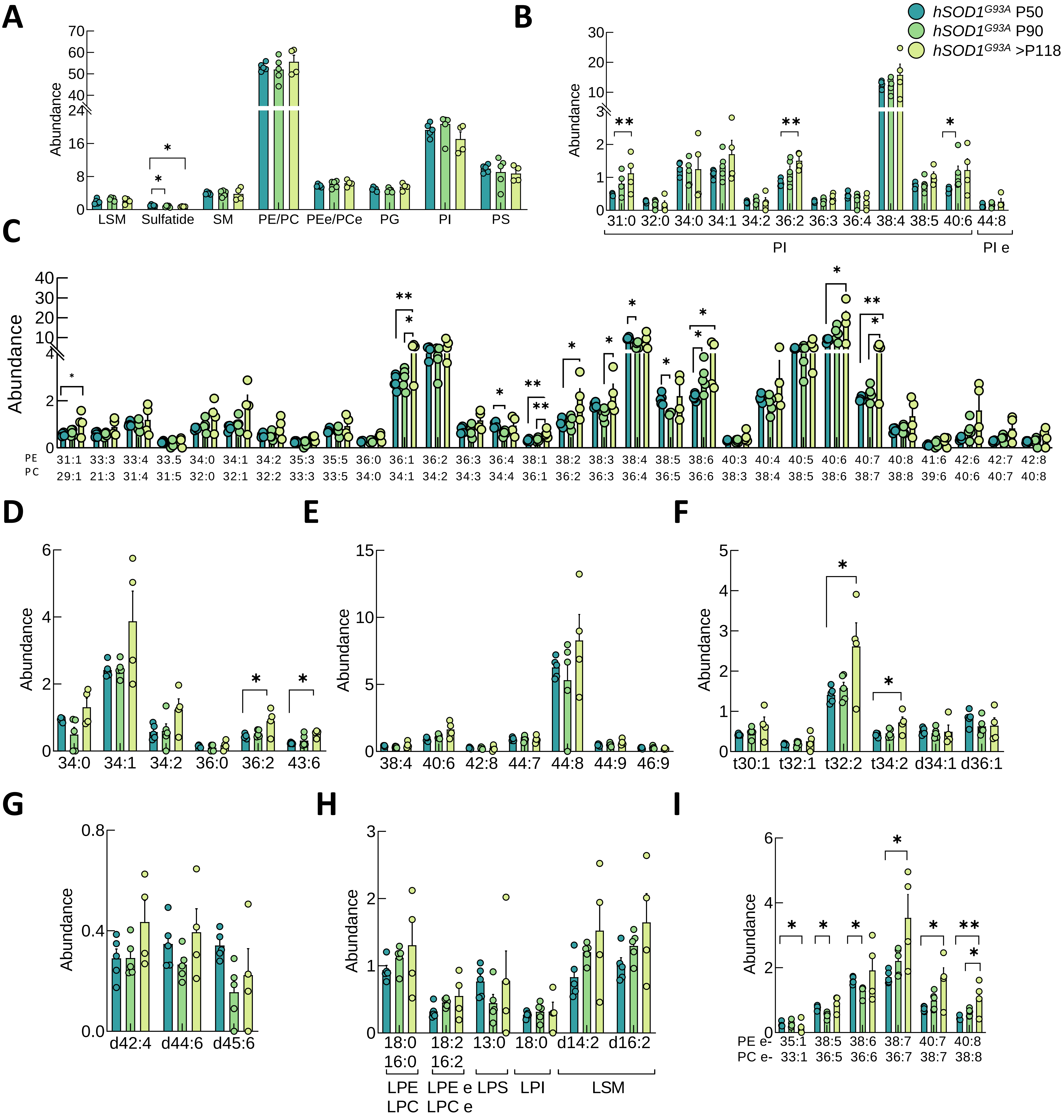


**Supplemental Figure 14.** Relative abundance of (**A**) different lipid families detected, and different lipid species of (**B**) PI and PIe, (**C**) PE/PC, (**D**) PG, (**E**) PS, (**F**) SM, (**G**) SFT, (**H**) LPE, LPC, LPEe, LPCe, LPS, LPI and LSM, and (**I**) PEe and PCe in *Tibialis anterior* muscle from IIX fibers of *hSOD1*$^{G93A}$ female mice at postnatal day 50 (presymptomatic), 90 (symptomatic) and > 118 (end-stage). Abbreviations: PI, phosphatidylinositols; PIe, ether-linked phosphatidylinositols; PE, phosphatidylethanolamines; PC, phosphatidylcholines; PG, phosphatidylglycerols; PS, phosphatidylserines; SM, sphingomyelins; SFT, sulfatides; LPE, lysophosphatidylethanolamines; LPC, lysophosphatidylcholines; LPEe, ether-linked lysophosphatidylethanolamines; LPCe, ether-linked lysophosphatidylcholines; LPI, lysophosphatidylinositols; LPS, lysophosphatidylserines; LSM, lysosphingomyelins; PEe, ether-linked phosphatidylethanolamines; PCe, ether-linked phosphatidylcholines. Bar graphs represent mean ± SEM. N≥4 per condition. For normally distributed data with equal variances, one-way ANOVA followed by Tukey's HSD post hoc test was applied. When variances were unequal, Welch ANOVA followed by Games-Howell post hoc test was used. For non-normally distributed data, the Kruskal-Wallis test followed Wilcoxon rank-sum test was applied. Significant differences are indicated as *$p$<0.05, **$p$<0.01, ***$p$<0.001.

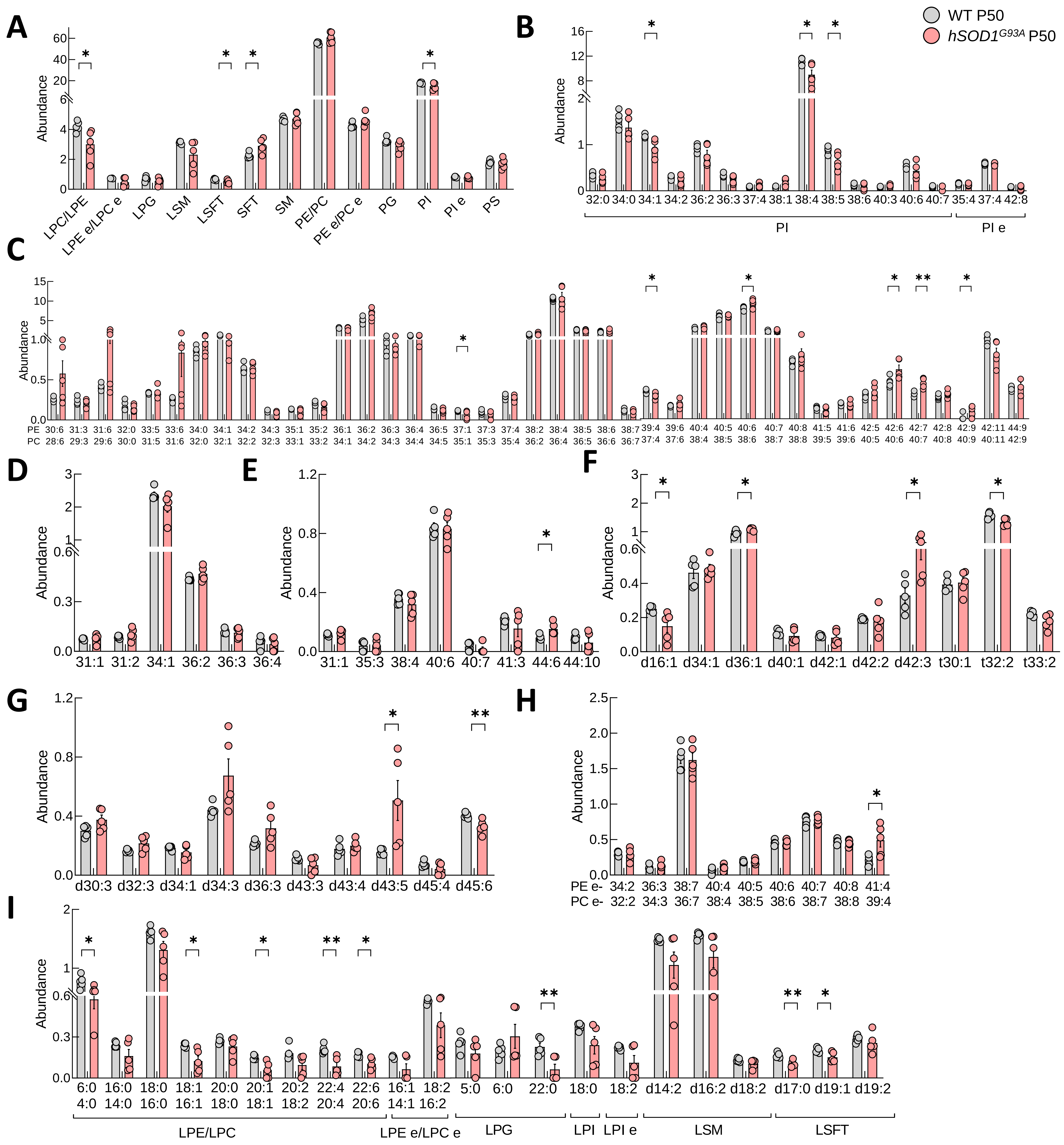


**Supplemental Figure 15.** Relative abundance of (**A**) different lipid families detected, and different lipid species of (**B**) PI and PIe, (**C**) PE/PC, (**D**) PG, (**E**) PS, (**F**) SM, (**G**) SFT, (**H**) PEe and Pce, and (**I**) LPE, LPC, LPEe, LPCe, LPG, LPI, LPIe, LSM and lysosulfatide in *Tibialis anterior* muscle of wild-type and *hSOD1*$^{G93A}$ male mice at postnatal day 50 (presymptomatic). Abbreviations: PI, phosphatidylinositols; PIe, ether-linked phosphatidylinositols; PE, phosphatidylethanolamines; PC, phosphatidylcholines; PG, phosphatidylglycerols; PS, phosphatidylserines; SM, sphingomyelins; SFT, sulfatides; LPE, lysophosphatidylethanolamines; LPC, lysophosphatidylcholines; LPEe, ether-linked lysophosphatidylethanolamines; LPCe, ether-linked lysophosphatidylcholines; LPG, lysophosphatidylglycerols; LPI, lysophosphatidylinositols; LPIe, ether-linked lysophosphatidylinositols; LSM, lysosphingomyelins; LSFT, lysosulfatides. Bar graphs represent mean ± SEM. N=5 per condition. For normally distributed data with equal variances, one-way ANOVA followed by Tukey's HSD post hoc test was applied. When variances were unequal, Welch ANOVA followed by Games-Howell post hoc test was used. For non-normally distributed data, the Kruskal-Wallis test followed Wilcoxon rank-sum test was applied. Significant differences are indicated as *$p$<0.05, **$p$<0.01, ***$p$<0.001.

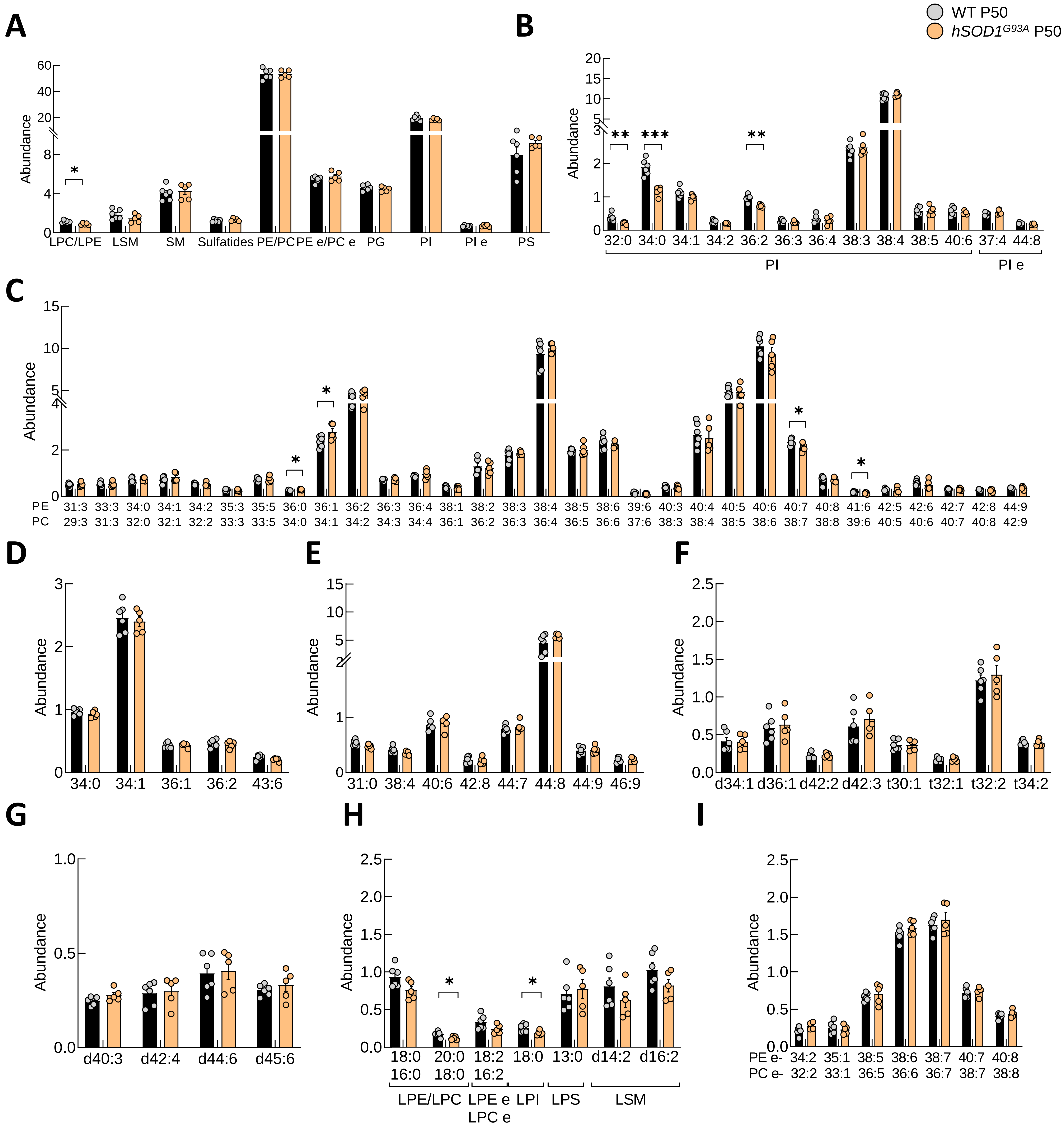


**Supplemental Figure 16.** Relative abundance of (**A**) different lipid families detected, and different lipid species of (**B**) PI and PIe, (**C**) PE/PC, (**D**) PG, (**E**) PS, (**F**) SM, (**G**) SFT, (**H**) LPE, LPC, LPEe, LPCe, LPI, LPS, and LSM and, (**I**) PEe and PCe in *Tibialis anterior* muscle from IIA fibers of wild-type and *hSOD1*$^{G93A}$ female mice at postnatal day 50 (presymptomatic). Abbreviations: PI, phosphatidylinositols; PIe, ether-linked phosphatidylinositols; PE, phosphatidylethanolamines; PC, phosphatidylcholines; PG, phosphatidylglycerols; PS, phosphatidylserines; SM, sphingomyelins; SFT, sulfatides; LPE, lysophosphatidylethanolamines; LPC, lysophosphatidylcholines; LPEe, ether-linked lysophosphatidylethanolamines; LPCe, ether-linked lysophosphatidylcholines; LPS, lysophosphatidylserines; LPI, lysophosphatidylinositols; LSM, lysosphingomyelins; PEe, ether-linked phosphatidylethanolamines; PCe, ether-linked phosphatidylcholines. Bar graphs represent mean ± SEM. N≥5 per condition. For normally distributed data with equal variances, one-way ANOVA followed by Tukey's HSD post hoc test was applied. When variances were unequal, Welch ANOVA followed by Games-Howell post hoc test was used. For non-normally distributed data, the Kruskal-Wallis test followed Wilcoxon rank-sum test was applied. Significant differences are indicated as *$p$<0.05, **$p$<0.01, ***$p$<0.001.

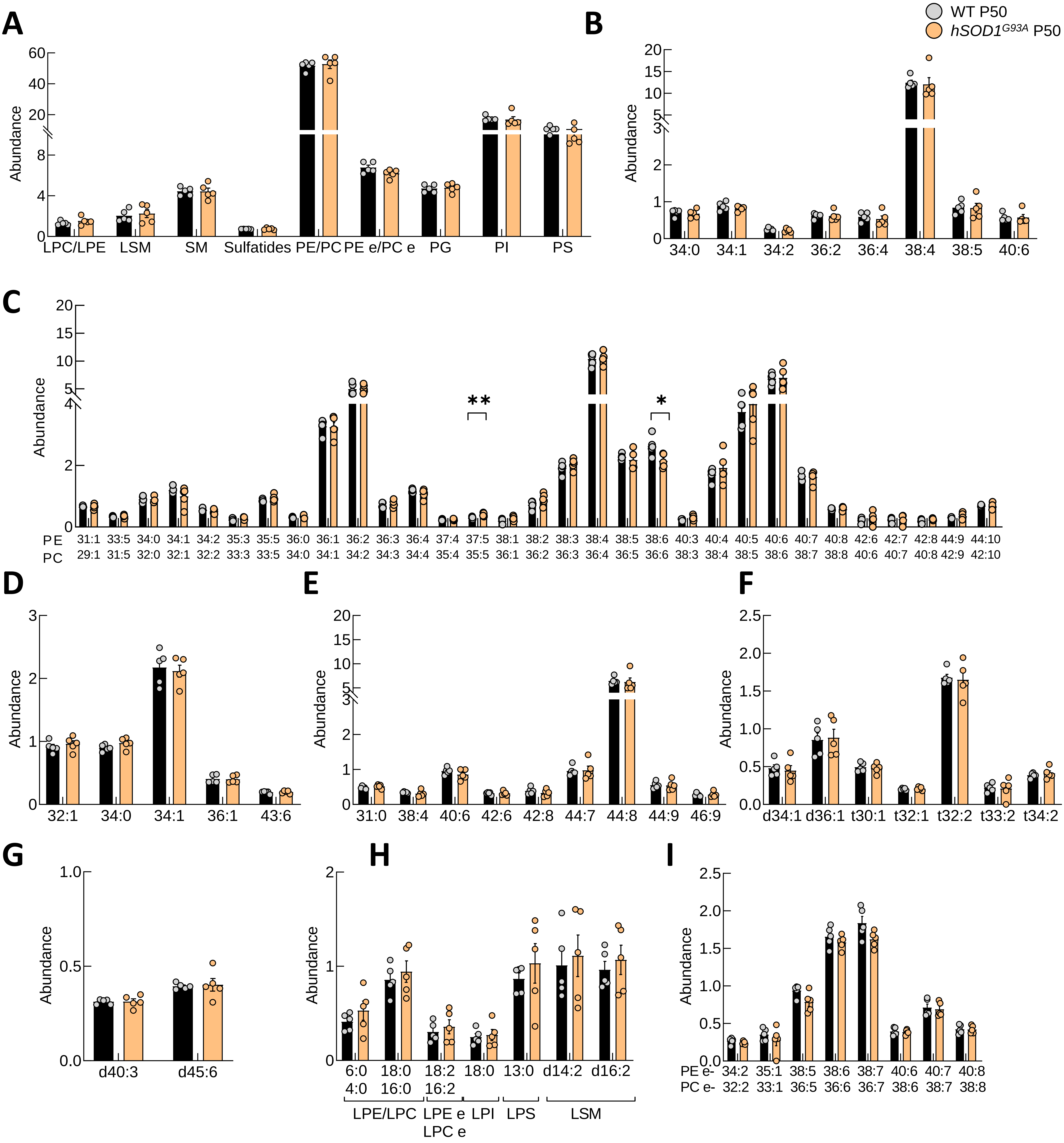


**Supplemental Figure 17.** Relative abundance of (**A**) different lipid families detected, and different lipid species of (**B**) PI, (**C**) PE/PC, (**D**) PG, (**E**) PS, (**F**) SM, (**G**) SFT, (**H**) LPE, LPC, LPEe, LPCe, LPI, LPS, and LSM and, (**I**) PEe and PCe in *Tibialis anterior* muscle from IIB fibers of wild-type and *hSOD1*$^{G93A}$ female mice at postnatal day 50 (presymptomatic). Abbreviations: PI, phosphatidylinositols; PE, phosphatidylethanolamines; PC, phosphatidylcholines; PG, phosphatidylglycerols; PS, phosphatidylserines; SM, sphingomyelins; SFT, sulfatides; LPE, lysophosphatidylethanolamines; LPC, lysophosphatidylcholines; LPEe, ether-linked lysophosphatidylethanolamines; LPCe, ether-linked lysophosphatidylcholines; LPS, lysophosphatidylserines; LPI, lysophosphatidylinositols; LSM, lysosphingomyelins; PEe, ether-linked phosphatidylethanolamines; PCe, ether-linked phosphatidylcholines. Bar graphs represent mean ± SEM. N≥5 per condition. For normally distributed data with equal variances, one-way ANOVA followed by Tukey's HSD post hoc test was applied. When variances were unequal, Welch ANOVA followed by Games-Howell post hoc test was used. For non-normally distributed data, the Kruskal-Wallis test followed Wilcoxon rank-sum test was applied. Significant differences are indicated as *$p$<0.05, **$p$<0.01, ***$p$<0.001.

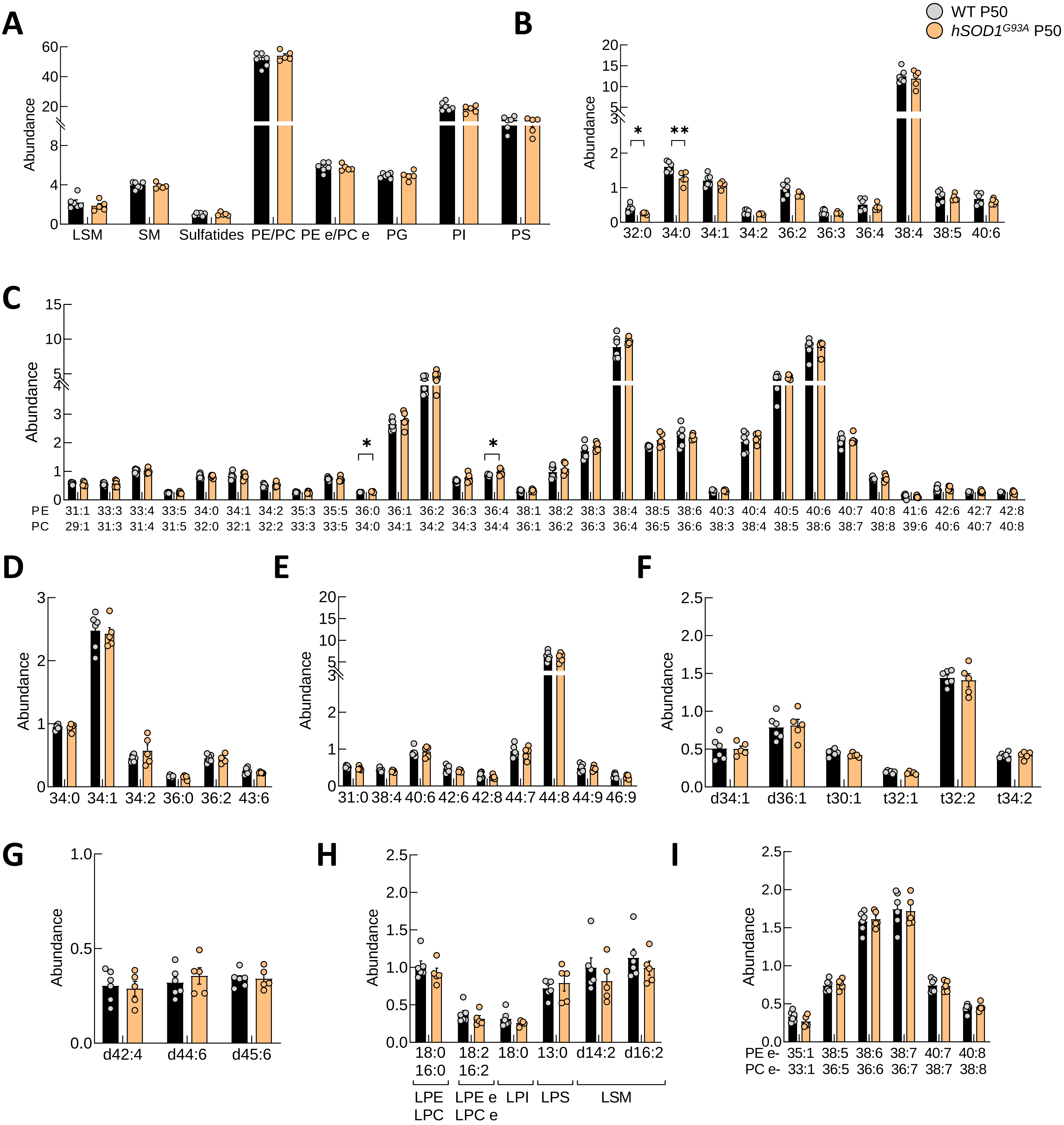


**Supplemental Figure 18.** Relative abundance of (**A**) different lipid families detected, and different lipid species of (**B**) PI, (**C**) PE/PC, (**D**) PG, (**E**) PS, (**F**) SM, (**G**) SFT, (**H**) LPE, LPC, LPEe, LPCe, LPI, LPS, and LSM and, (**I**) PEe and PCe in *Tibialis anterior* muscle from IIX fibers of wild-type and *hSOD1*$^{G93A}$ female mice at postnatal day 50 (presymptomatic). Abbreviations: PI, phosphatidylinositosl; PE, phosphatidylethanolamines; PC, phosphatidylcholines; PG, phosphatidylglycerols; PS, phosphatidylserines; SM, sphingomyelins; SFT, sulfatides; LPE, lysophosphatidylethanolamines; LPC, lysophosphatidylcholines; LPEe, ether-linked lysophosphatidylethanolamines; LPCe, ether-linked lysophosphatidylcholines; LPS, lysophosphatidylserines; LPI, lysophosphatidylinositols; LSM, lysosphingomyelins; PEe, ether-linked phosphatidylethanolamines; PCe, ether-linked phosphatidylcholines. Bar graphs represent mean ± SEM. N≥5 per condition. For normally distributed data with equal variances, one-way ANOVA followed by Tukey's HSD post hoc test was applied. When variances were unequal, Welch ANOVA followed by Games-Howell post hoc test was used. For non-normally distributed data, the Kruskal-Wallis test followed Wilcoxon rank-sum test was applied. Significant differences are indicated as *$p$<0.05, **$p$<0.01, ***$p$<0.001.

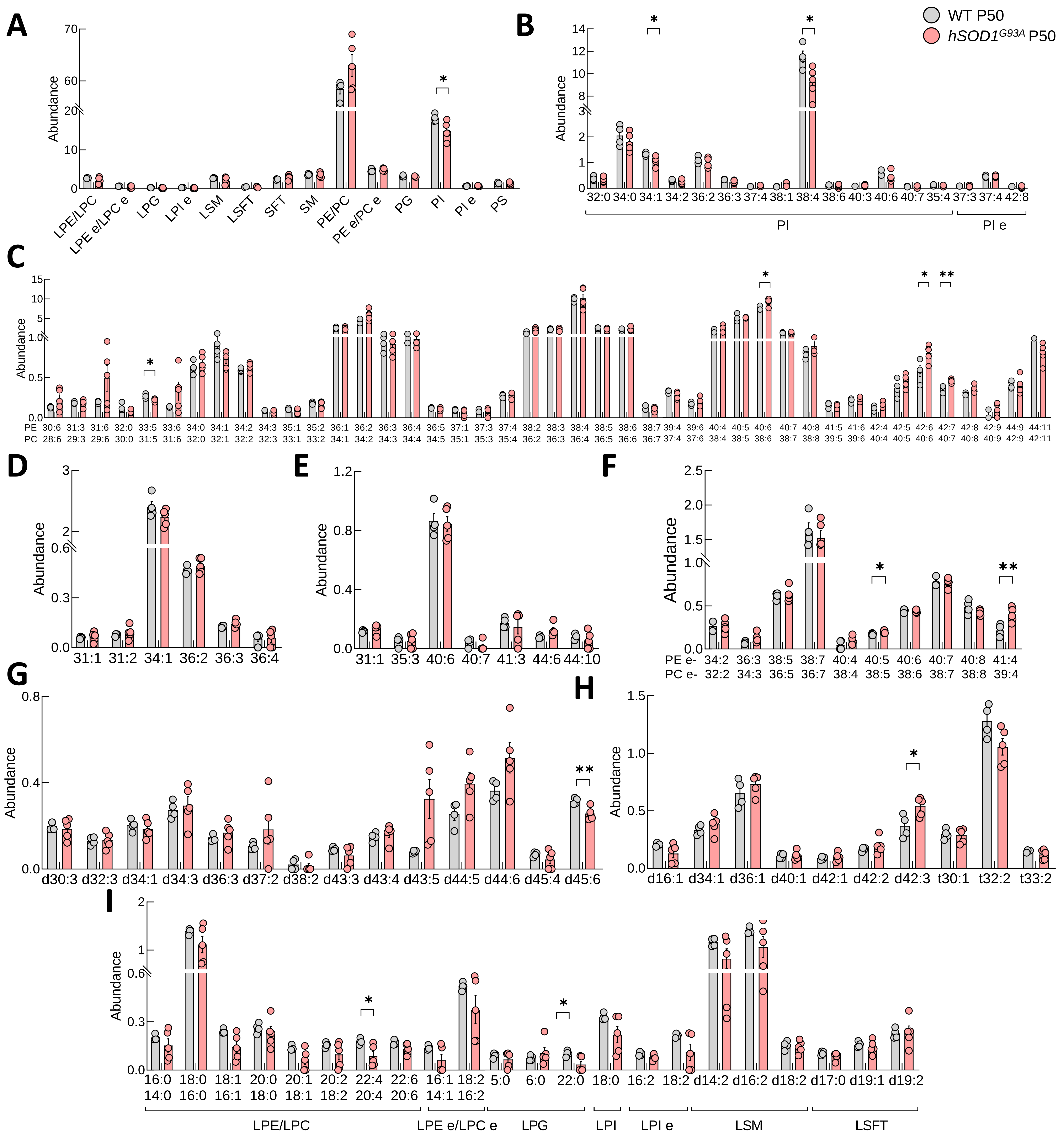


**Supplemental Figure 19.** Relative abundance of (**A**) different lipid families detected, and different lipid species of (**B**) PI and PIe, (**C**) PE/PC, (**D**) PG, (**E**) PS, (**F**) PEe and PCe, (**G**) SFT, (**H**) SM and, (**I**) LPE, LPC, LPEe, LPCe, LPG, LPI, LPIe, LSM and lysosulfatide in *Tibialis anterior* muscle from IIA fibers of wild-type and *hSOD1$^{G93A}$* male mice at postnatal day 50 (presymptomatic). Abbreviations: PI, phosphatidylinositols; PIe, ether-linked phosphatidylinositols; PE, phosphatidylethanolamines; PC, phosphatidylcholines; PG, phosphatidylglycerols; PS, phosphatidylserines; SM, sphingomyelins; SFT, sulfatides; LPE, lysophosphatidylethanolamines; LPC, lysophosphatidylcholines; LPEe, ether-linked lysophosphatidylethanolamines; LPCe, ether-linked lysophosphatidylcholines; LPG, lysophosphatidylglycerols; LPI, lysophosphatidylinositols; LPIe, ether-linked lysophosphatidylinositols; LSM, lysosphingomyelins; LSFT, lysosulfatides. Bar graphs represent mean ± SEM. N=5 per condition. For normally distributed data with equal variances, one-way ANOVA followed by Tukey's HSD post hoc test was applied. When variances were unequal, Welch ANOVA followed by Games-Howell post hoc test was used. For non-normally distributed data, the Kruskal-Wallis test followed Wilcoxon rank-sum test was applied. Significant differences are indicated as *$p$<0.05, **$p$<0.01, ***$p$<0.001.

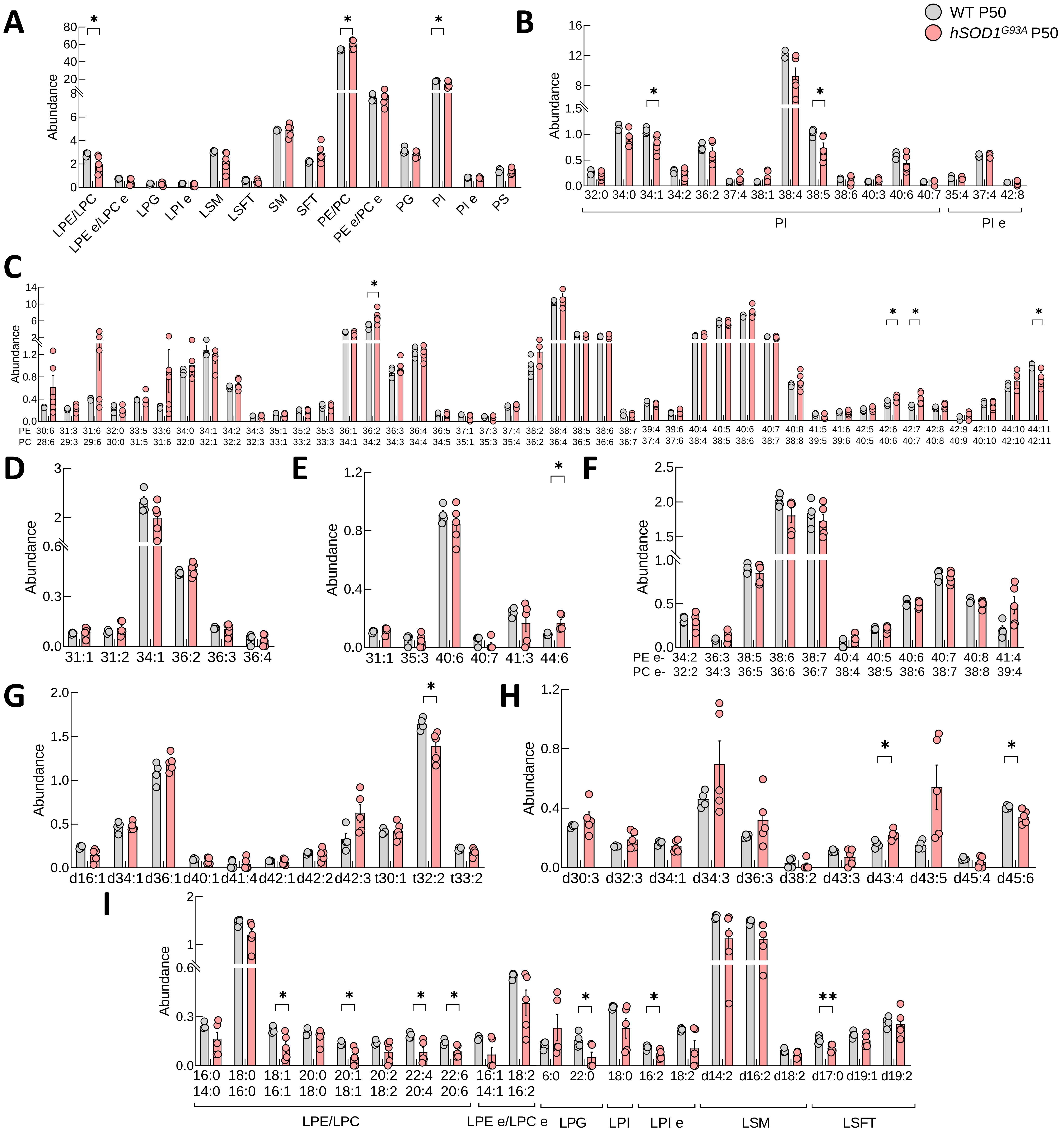


**Supplemental Figure 20.** Relative abundance of (**A**) different lipid families detected, and different lipid species of (**B**) PI and PIe, (**C**) PE/PC, (**D**) PG, (**E**) PS, (**F**) PEe and PCe, (**G**) SM, (**H**) SFT and, (**I**) LPE, LPC, LPEe, LPCe, LPG, LPI, LPIe, LSM and lysosulfatide in *Tibialis anterior* muscle from IIB fibers of wild-type and *hSOD1*$^{G93A}$ male mice at postnatal day 50 (presymptomatic). Abbreviations: PI, phosphatidylinositols; PIe, ether-linked phosphatidylinositols; PE, phosphatidylethanolamines; PC, phosphatidylcholines; PG, phosphatidylglycerols; PS, phosphatidylserines; SM, sphingomyelins; SFT, sulfatides; LPE, lysophosphatidylethanolamines; LPC, lysophosphatidylcholines; LPEe, ether-linked lysophosphatidylethanolamines; LPCe, ether-linked lysophosphatidylcholines; LPG, lysophosphatidylglycerols; LPI, lysophosphatidylinositols; LPIe, ether-linked lysophosphatidylinositols; LSM, lysosphingomyelins; LSFT, lysosulfatides. Bar graphs represent mean ± SEM. N≥4 per condition. For normally distributed data with equal variances, one-way ANOVA followed by Tukey's HSD post hoc test was applied. When variances were unequal, Welch ANOVA followed by Games-Howell post hoc test was used. For non-normally distributed data, the Kruskal-Wallis test followed Wilcoxon rank-sum test was applied. Significant differences are indicated as *$p$<0.05, **$p$<0.01, ***$p$<0.001.

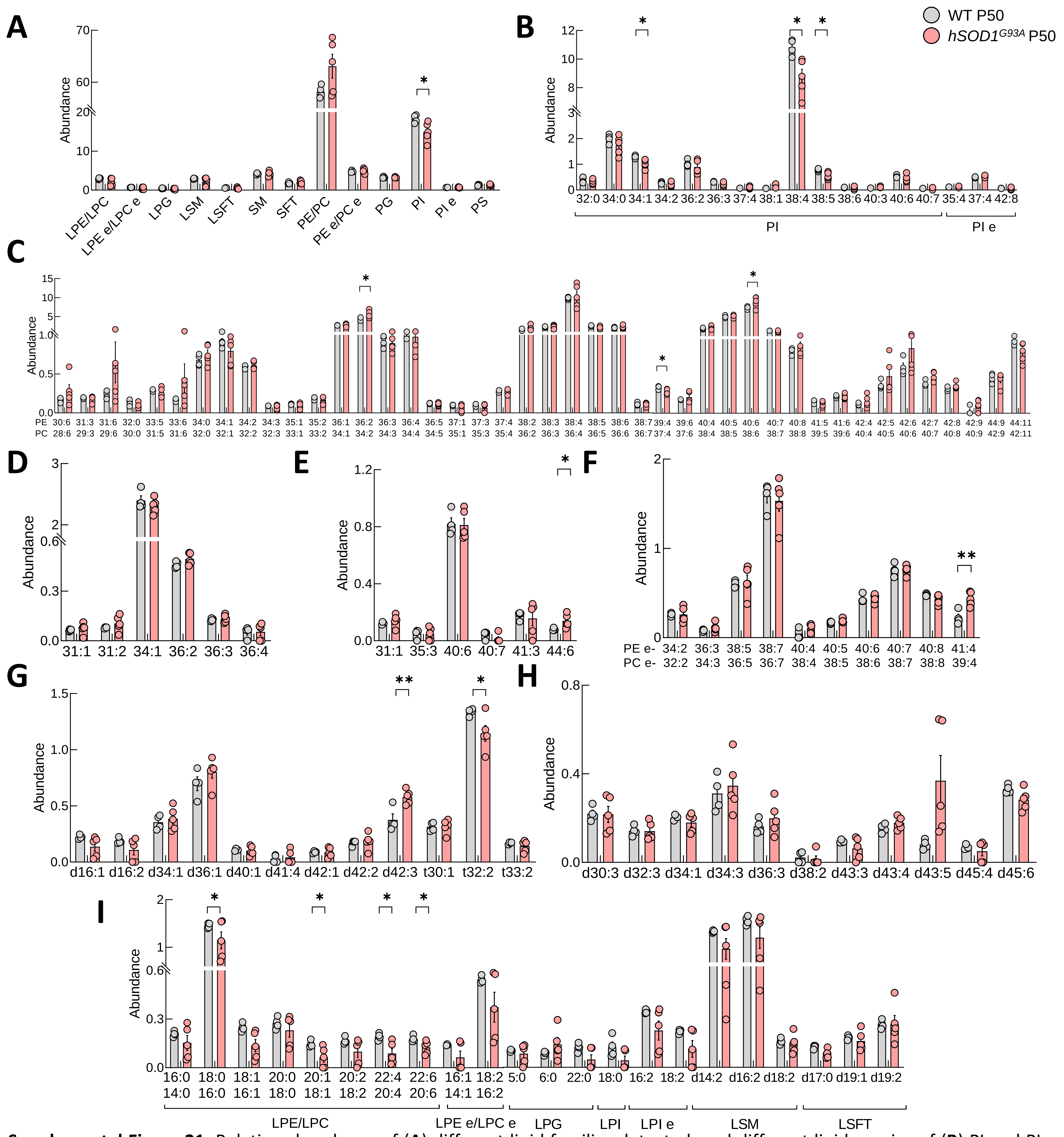


**Supplemental Figure 21.** Relative abundance of (**A**) different lipid families detected, and different lipid species of (**B**) PI and PIe, (**C**) PE/PC, (**D**) PG, (**E**) PS, (**F**) PEe and PCe, (**G**) SFT, (**H**) SM and, (**I**) LPE, LPC, LPEe, LPCe, LPG, LPI, LPIe, LSM and lysosulfatide in *Tibialis anterior* muscle from IIX fibers of wild-type and *hSOD1*$^{G93A}$ male mice at postnatal day 50 (presymptomatic). Abbreviations: PI, phosphatidylinositols; PIe, ether-linked phosphatidylinositols; PE, phosphatidylethanolamines; PC, phosphatidylcholines; PG, phosphatidylglycerols; PS, phosphatidylserines; SM, sphingomyelins; SFT, sulfatides; LPE, lysophosphatidylethanolamines; LPC, lysophosphatidylcholines; LPEe, ether-linked lysophosphatidylethanolamines; LPCe, ether-linked lysophosphatidylcholines; LPG, lysophosphatidylglycerols; LPI, lysophosphatidylinositols; LPIe, ether-linked lysophosphatidylinositols; LSM, lysosphingomyelins; LSFT, lysosulfatides. Bar graphs represent mean ± SEM. N≥4 per condition. For normally distributed data with equal variances, one-way ANOVA followed by Tukey's HSD post hoc test was applied. When variances were unequal, Welch ANOVA followed by Games-Howell post hoc test was used. For non-normally distributed data, the Kruskal-Wallis test followed Wilcoxon rank-sum test was applied. Significant differences are indicated as *$p$<0.05, **$p$<0.01, ***$p$<0.001.
